# Automated quality control and cell identification of droplet-based single-cell data using dropkick

**DOI:** 10.1101/2020.10.08.332288

**Authors:** Cody N. Heiser, Victoria M. Wang, Bob Chen, Jacob J. Hughey, Ken S. Lau

**Affiliations:** Epithelial Biology Center, Vanderbilt University Medical Center, 2213 Garland Avenue, 10475 MRB IV, Nashville, TN 37232, USA; Program in Chemical and Physical Biology, Vanderbilt University School of Medicine, Nashville, TN 37232, USA; Department of Computer Science, Vanderbilt University, Nashville, TN 37232, USA; Department of Biomedical Informatics, Vanderbilt University School of Medicine, Nashville, TN 37232, USA; Department of Biological Sciences, Vanderbilt University, Nashville, TN 37232, USA; Department of Cell and Developmental Biology, Vanderbilt University School of Medicine, Nashville, TN 37232, USA; Center for Quantitative Sciences, Vanderbilt University School of Medicine, Nashville, TN 37232, USA

## Abstract

A major challenge for droplet-based single-cell sequencing technologies is distinguishing true cells from uninformative barcodes in datasets with disparate library sizes confounded by high technical noise (i.e. batch-specific ambient RNA). We present dropkick, a fully automated software tool for quality control and filtering of single-cell RNA sequencing (scRNA-seq) data with a focus on excluding ambient barcodes and recovering real cells bordering the quality threshold. By automatically determining dataset-specific training labels based on predictive global heuristics, dropkick learns a gene-based representation of real cells and ambient noise, calculating a cell probability score for each barcode. Using simulated and real-world scRNA-seq data, we benchmarked dropkick against a conventional thresholding approach and EmptyDrops, a popular computational method, demonstrating greater recovery of rare cell types and exclusion of empty droplets and noisy, uninformative barcodes. We show for both low and high-background datasets that dropkick’s weakly supervised model reliably learns which genes are enriched in ambient barcodes and draws a multidimensional boundary that is more robust to dataset-specific variation than existing filtering approaches. dropkick provides a fast, automated tool for reproducible cell identification from scRNA-seq data that is critical to downstream analysis and compatible with popular single-cell analysis Python packages.

## Introduction

Single-cell RNA sequencing (scRNA-seq) allows for untargeted profiling of genome-scale expression in thousands of individual cells, providing insights into tissue heterogeneity and population dynamics. Droplet-based platforms that involve microfluidic encapsulation of cells in water-oil emulsions (Klein, et al. 2015; Macosko, et al. 2015; Zheng, et al. 2017) have grown widely popular for their robustness and throughput. The use of barcoded poly-thymidine capture oligonucleotides provides information for assigning eventual sequencing reads to each droplet downstream of bulk library preparation. Due to the low cellular density required to avoid doublets (i.e., two or more cells captured in the same droplet), the vast majority of droplets are empty, ideally containing only tissue dissociation buffer and a barcoded RNA-capture bead with no cellular RNA. However, during the tissue dissociation process, cell death, lysis, and leakage result in the shedding of ambient mRNA into the supernatant solution, which is then captured as background in droplets containing individual cells and so-called “empty droplet” reactions. Ultimately, a droplet-based scRNA-seq dataset contains up to hundreds of thousands of barcodes that correspond to these “empty droplets” which include sequenced material from ambient RNA alone.

In order to prepare these data for downstream analysis, empty droplets and other “junk” barcodes with little to no molecular information must be removed. Often, computational biologists will define manual thresholds on global heuristics such as total counts of unique molecular identifiers (UMI) or the total number of genes detected in each barcode in order to isolate high-quality cells. While these hard cutoffs may generally yield expected cell populations and remove the bulk of populational noise in low-background samples, they are highly arbitrary, batch-specific, and generally biased against cell types with low RNA content or genetic diversity (Lun, et al. 2019). Furthermore, lenient thresholds often yield filtered datasets with populations of dead and dying cells or empty droplets with high ambient RNA content, especially in encapsulations with high background resulting from tissue-specific cell viability and dissociation protocols. These cell clusters may be gated out manually by the experienced single-cell biologist, but they will distort dimension-reduced embeddings and alter statistical testing for differential gene expression if left unchecked.

Here we introduce dropkick, a fully automated machine learning software tool for data-driven filtering of droplet-based scRNA-seq data. dropkick provides a quality control (QC) module for initial evaluation of global distributions that define barcode populations (real cells vs. empty droplets) and quantifies the batch-specific ambient gene profile. The dropkick filtering module establishes initial thresholds on predictive global heuristics using an automated gradient-descent method, then trains a gene-based logistic regression model to assign confidence scores to all barcodes in the dataset. dropkick model coefficients are sparse and biologically informative, identifying a minimal number of gene features associated with empty droplets and low-quality cells in a weakly supervised fashion. We show that dropkick outperforms basic threshold-based filtering and a similar data-driven model (Lun, et al. 2019) in recovery of expected cell types and exclusion of empty droplets, with robustness and reproducibility across encapsulation platforms, samples, and varying degrees of noise from ambient RNA.

## Results

### Evaluating dataset quality with the dropkick QC module

Global data quality and predominance of ambient RNA affect both reliable cell identification as well as downstream analyses including clustering, cell type annotation, and trajectory inference in scRNA-seq data (Young and Behjati 2018; Fleming, et al. 2019; Yang, et al. 2020). Single-cell data with a low signal-to-noise ratio due to high ambient background can result in information loss that may ultimately confound cell type and cell state identification and related statistical analyses (Zhang, et al. 2019). For instance, a scRNA-seq encapsulation with a high degree of cell lysis can cause highly expressed marker genes from abundant cell types to be present in the ambient RNA profile that contaminates all cell barcodes. In this scenario, global differences between cell populations would be diminished by the common detection of ambient noise, leading to loss of resolution in inference of cell identity and state.

In order to quantify ambient contamination that reduces this batch-specific signal-to-noise ratio, we have developed a comprehensive quality control report for unfiltered, post-alignment UMI counts matrices. Figure 1 provides an example dropkick QC report for a human T cell dataset encapsulated using the 10X Genomics Chromium platform (Zheng, et al. 2017). This sample is exemplary of a low-background dataset, as the cells isolated from human blood do not require dissociation that causes cell stress and lysis in other tissues (Supplementary Figure 1). Barcodes are ranked by total counts to yield a profile that describes the expected number of high-quality cells, empty droplets, and “junk” barcodes (Figure 1A; Fleming et al. 2019). The number of genes detected per barcode follows a similar distribution to total counts, which informs our choice of dropkick training thresholds in the following sections. The first plateau in the total counts profile of the T cell dataset indicates approximately 4,000 high-quality cells, followed by a sharp drop in the distribution (Figure 1A). This drop-off in total UMI content signifies an estimated location for a manual cutoff as seen in the 10X CellRanger version 2 analysis software (Lun, et al. 2019).

**Figure 1.**
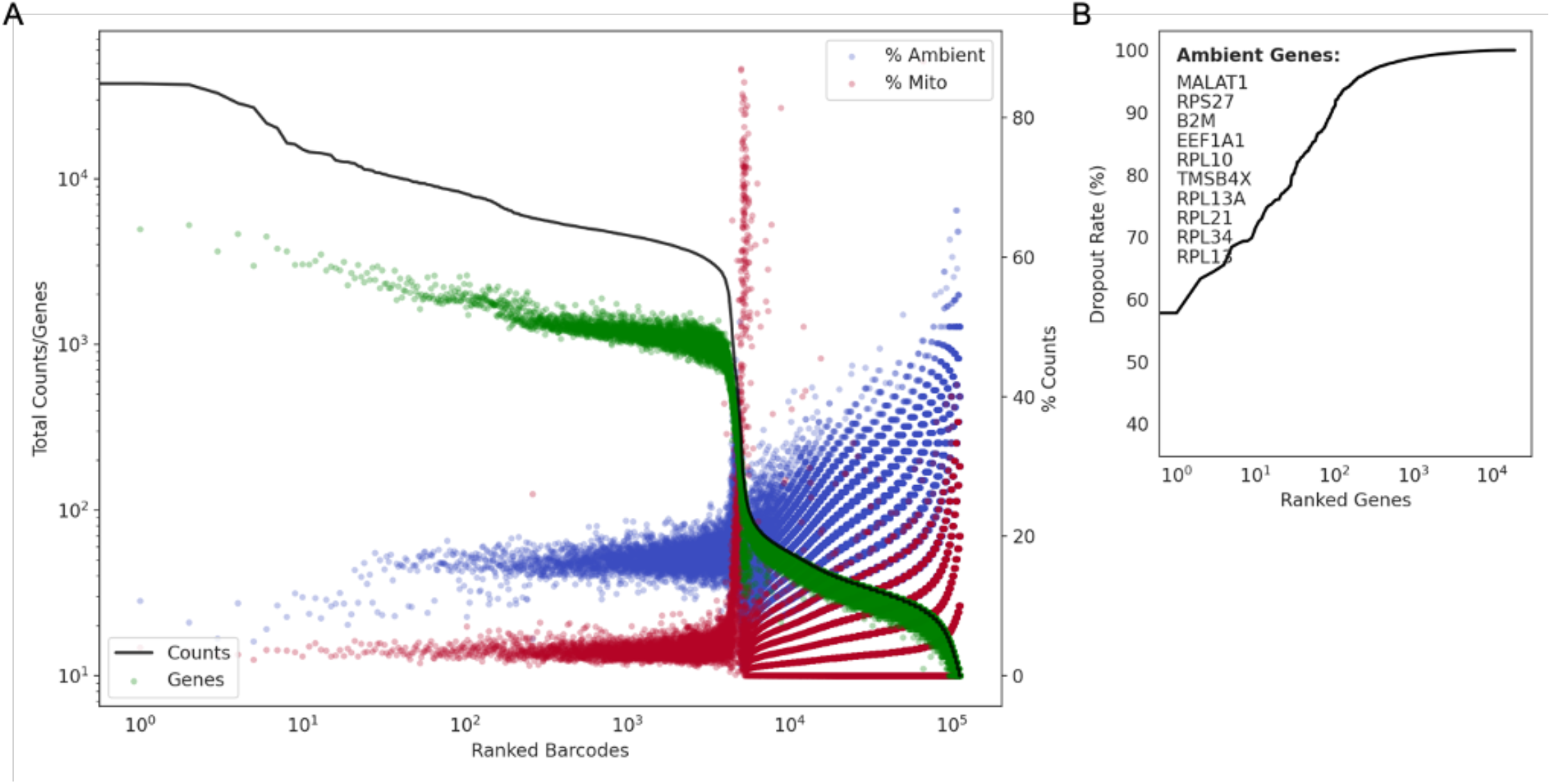
Evaluating dataset quality with the dropkick QC module. A) Profile of total counts (black trace) and genes (green points) detected per ranked barcode in the 4k pan-T cell dataset (10X Genomics). Percentage of mitochondrial (red) and ambient (blue) reads for each barcode included to denote quality along dataset profile. B) Profile of dropout rate per ranked gene. Ambient genes are identified by dropkick and used to calculate ambient percentage in A.

dropkick next defines a subset of ambient genes using the dropout rate, or the fraction of barcodes in which each gene is not detected. Ranking genes from lowest to highest dropout rate (Figure 1B), dropkick labels those with dropout rates lower than the top ten as “ambient”. High-background datasets may have many genes that are detected in every barcode (dropout rate = 0). The dropkick definition of an ambient profile thus ensures that all relevant genes are included. The contribution of this ambient subset to the total counts of each barcode can then be calculated, shown as blue points in the dropkick QC report (Figure 1A). Similarly, an overlay of mitochondrial read percentage indicates dead or dying cells undergoing apoptosis (Tait and Green 2010). Indeed, the ambient and mitochondrial contributions to the empty droplets in the second plateau of the total counts log-rank curve is markedly higher than the first plateau (Figure 1A). Another noteworthy observation is dropkick defines an ambient profile that is distinct from the subset of mitochondrial genes. This is important for assessing cell quality in downstream clustering and dimension reduction, as any empty droplets that remain in the dataset post-filtering often cluster together in low-dimensional embeddings and can be highlighted by their enrichment in ambient genes. As stated previously, marker genes from abundant cell types may show up in the ambient gene set due to excessive lysis of these common cells during tissue preparation (Young and Behjati 2018; Fleming, et al. 2019; Yang, et al. 2020; Supplementary Figure 1). Accordingly, analysts should be cognizant of background expression levels that contaminate adjacent cell populations and confound cell type identification during subsequent analysis.

As each scRNA-seq dataset has unique, batch-specific ambient RNA profiles and barcode distributions, the dropkick QC module allows for estimation of global data quality. Mouse colonic mucosa dissociated and encapsulated in parallel using inDrop and 10X Genomics platforms (Supplementary Figure 1) exemplifies high-background scRNA-seq data, as indicated by elevated RNA levels in the second plateau of the total counts and genes curves. Moreover, marker genes *Car1* and *Muc2* from abundant colonocytes and goblet cells, respectively, are identified by dropkick as ambient genes for these datasets. This signifies lysis of common epithelial cell populations during tissue preparation and dissociation. Given the dropkick QC report, the user should thus expect background expression across all barcodes, which could prove pivotal to downstream processing and biological interpretation. Taken together, dropkick can estimate the number of high-quality cells in our dataset, determine average background noise from ambient RNA, and thus predict performance of filtering and ensuing analysis based on global data quality.

### Description of dropkick filtering method

dropkick uses weakly supervised machine learning to build a model of single-cell gene expression in order to score and classify barcodes as real cells or empty droplets within individual scRNA-seq datasets. To construct a training set for this model, dropkick begins by calculating batch-specific global metrics that are generally predictive of barcode quality, such as the total number of genes detected (n_genes; Figure 2A) which was chosen as the default training heuristic for dropkick by testing concordance with three alternative cell labels across 46 scRNA-seq samples (Supplementary Figure 2). A dataset similar to the 10X Genomics human T cell encapsulation (Figure 1A) will exhibit a multimodal distribution of n_genes across all barcodes (Figure 2B) where the peaks of the distribution match the plateaus seen in the log-rank representation (Figure 2C). Next, dropkick performs multi-level thresholding on the n_genes histogram using Otsu’s method (Otsu 1979; Figure 2B,C). This automated gradient-descent technique divides the barcode distribution into three levels in this “heuristic space”: a lower level containing uninformative “junk” barcodes (which are thrown away), an upper level containing barcodes with very high cell probability based on n_genes, and an intermediate level that consists of both high-RNA empty droplets and relatively low-RNA cells. The upper and intermediate barcode populations are labeled as real cells and putative empty droplets, respectively, for initial dropkick model training. These weakly self-supervised labels based on threshold cutoffs in “heuristic space” are expected to be noisy, and the goal of the next step of the dropkick pipeline is to re-draw these rough boundaries in “gene space” using logistic regression in order to recover real cells from the intermediate barcode cohort while removing ambient barcodes from the upper plateau (Figure 2D,E).

**Figure 2.**
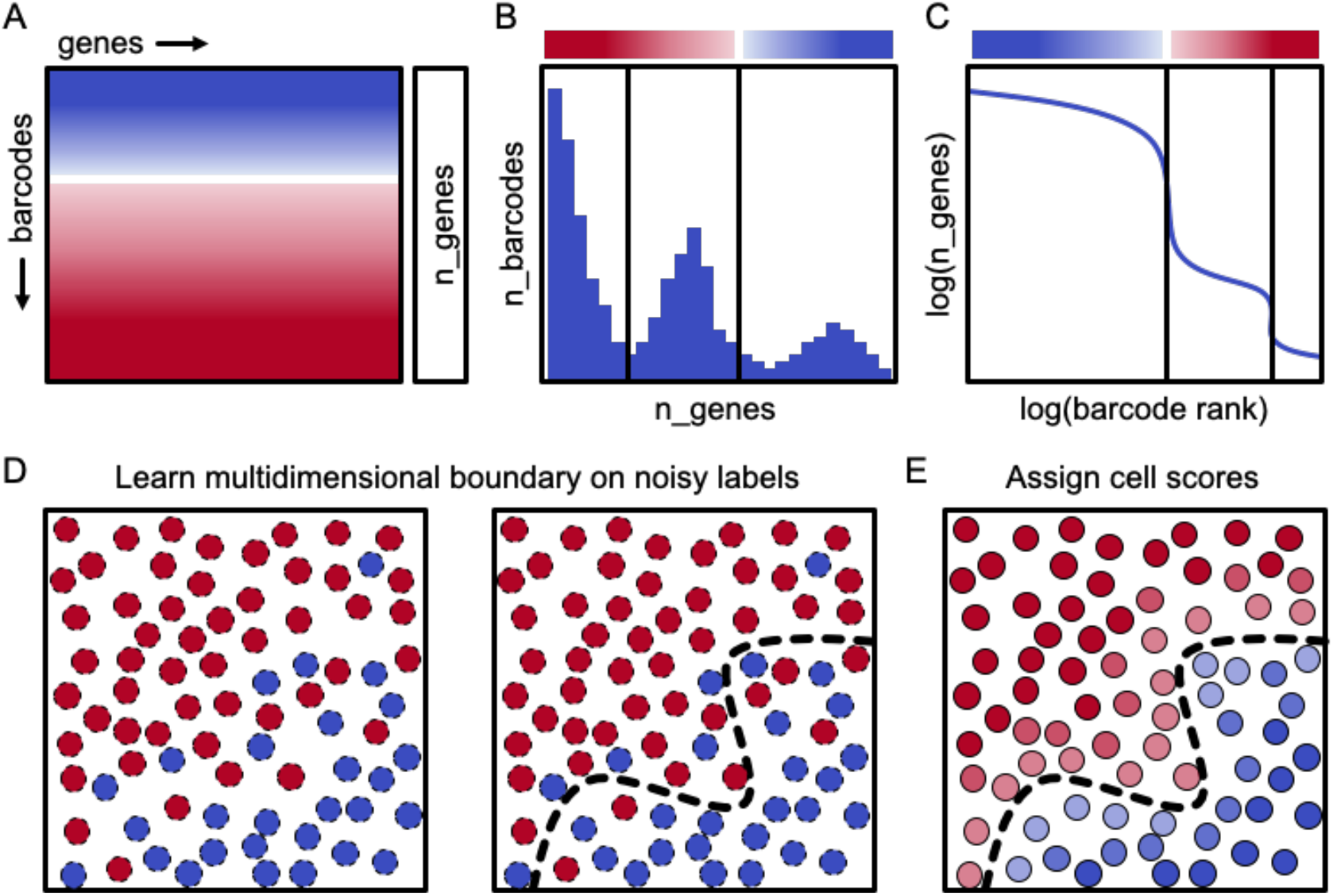
Description of dropkick filtering method. A) Diagram of scRNA-seq counts matrix with initial cell confidence for each barcode based solely on total genes detected (n_genes), depicted by color (red = empty droplet, blue = real cell). B) Histogram showing the distribution of barcodes by their n_genes value. Black lines indicate automated thresholds for training the dropkick model. C) log(n_genes) vs. log(rank) representation of barcode distribution as in dropkick QC report (Figure 1A). Thresholds from B are superimposed. D) Thresholds in heuristic space (B-C) are used to define initial training labels for logistic regression. E) dropkick chooses an optimal regularization strength through cross-validation, then assigns cell probabilities and labels to all barcodes using the trained model in gene space.

The logistic regression model employed by dropkick uses elastic net regularization (Zou and Hastie 2005), which balances feature selection and grouping by preserving or removing correlated genes from the model in concert. Ultimately, the motivation for choosing this regularization method is two-fold. First, the resulting model exists in “gene space”, maintaining the relative dimensionality of the dataset and providing biologically interpretable coefficients that describe barcode quality. Second, the model is penalized for complexity, which yields the simplest model (sparse coefficients) that adjusts the noisy initial labels while compensating for expected collinearities and errors in measurement.

### Evaluating dropkick filtering performance with simulated data

We tested dropkick filtering on single-cell data simulations that define both empty droplets and real cells, providing ground-truth labels for comparison to dropkick outputs (Fleming, et al. 2019). These simulations modeled ambient RNA noise in the cell populations to confound filtering, as seen in real-world datasets. We simulated both low (Figure 3A,B) and high (Figure 3C,D) background scenarios (see Methods: scRNA-seq data simulation).

**Figure 3.**
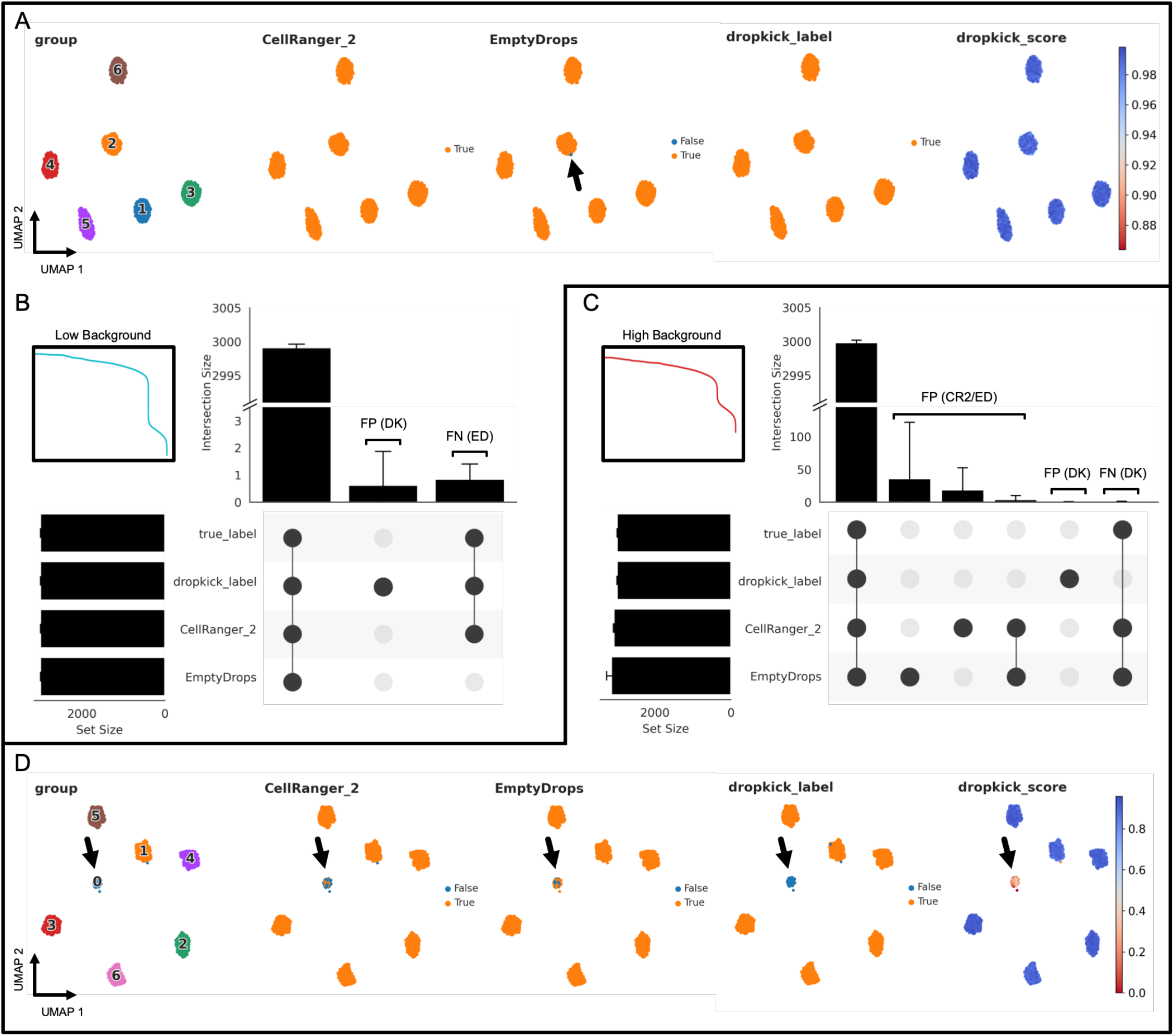
Evaluating dropkick filtering performance with simulated data. A) UMAP embedding of all barcodes kept by dropkick_label, CellRanger_2 and EmptyDrops for an example low-background simulation. Points colored by each of the three filtering labels, as well as ground-truth clusters determined by the simulation and dropkick score (cell probability). Arrow highlights a single false negative (FN) barcode in the EmptyDrops label set for this replicate. B) UpSet plot showing mean size of shared barcode sets across dropkick_label, CellRanger_2, EmptyDrops, and true labels for ten simulations. Error bars represent standard deviation. Unique sets show false positive (FP) barcodes labeled by dropkick and false negative (FN) barcodes excluded by EmptyDrops. Inset shows log-rank representation of the low-background simulation in A. C) Same as in B, for ten high-background simulations. Inset shows log-rank representation of the high-background simulation in D. D) Same as in A, for an example high-background simulation. Arrow highlights cluster 0, designated as “empty droplets” by simulation (see Methods: scRNA-seq data simulation).

To demonstrate the utility of the dropkick model over one-dimensional thresholding and an analogous data-driven filtering model, we ran dropkick, 10X Genomics CellRanger version 2 (CellRanger_2) and the EmptyDrops R package (Lun, et al. 2019) on ten iterations of low and high-background simulations. An example UMAP embedding of all barcodes kept by dropkick_label (dropkick score ≥ 0.5) and the two analogous methods shows that all three methods excluded empty droplets (assigned cluster 0 from the simulation), with a single false negative (FN) barcode highlighted in the EmptyDrops label set (Figure 3A). An UpSet plot (Figure 3B; Lex, et al. 2014) tabulating shared barcode sets across ten low-background simulations reveals nearly perfect specificity, sensitivity, and area under the receiver operating characteristic curve (AUROC) for all three methods in the low-background scenario (Supplementary Figure 3A,B,D, Supplementary Table 1, Supplementary Table 2).

Conversely, the high-background simulations produced a large number of false positives (FP) in the CellRanger_2 and EmptyDrops labels (Figure 3C), as ambient barcodes with high RNA content lie above the total counts threshold identified by CellRanger and the inflection point used as a testing cutoff by EmptyDrops (Lun, et al. 2019). A UMAP embedding of an example high-background simulation reveals a large population of empty droplets (assigned cluster 0 by the simulation) that dropkick_label removes from the final dataset (Figure 3D). Accordingly, dropkick displayed overall specificity and AUROC of 0.9999 ± 0.0002 and 0.9998 ± 0.0002 for the high-background simulations compared to 0.9910 ± 0.0018 and 0.9955 ± 0.0009 for CellRanger_2 and 0.9838 ± 0.0133 and 0.9917 ± 0.0071 for EmptyDrops, respectively (Supplementary Figure 3E,F,H, Supplementary Table 1, Supplementary Table 2).

We also compared outputs from the trained model (dropkick_label) to automated dropkick training labels (thresholding on n_genes) in both low- and high-background scenarios to further demonstrate the utility of dropkick’s machine learning model over heuristic cutoffs alone. Similar to CellRanger_2, the dropkick threshold performed favorably for the low background simulation, where real cells are separated distinctly from empty droplets in heuristic space – indicated by a sharp drop-off in total counts and genes in the dropkick QC log-rank plot (Figure 3B, inset). This one-dimensional thresholding resulted in sensitivity, specificity, and AUROC of 0.9986 ± 0.0007, 0.997 ± 0.0006, and 0.9978 ± 0.0005, respectively for ten low-background simulations (Supplementary Figure 3E, Supplementary Table 1). The trained dropkick model, on the other hand, recovered all real cells (sensitivity 1.0), with a perfect average AUROC of 1.0 ± 0.0 (Supplementary Figure 3D, Supplementary Table 1). This modest improvement indicates the utility of the dropkick model for sensitively discerning real cells from ambient barcodes over simple heuristic thresholding, even in a relatively low-background sample. In the high-background simulations, sensitivity of dropkick training labels fell to 0.8762 ± 0.0092 with an average AUROC of 0.9074 ± 0.0043 (Supplementary Figure 3G, Supplementary Table 1). Following model training, dropkick’s sensitivity and AUROC once again improved to 0.9995 ± 0.0004 and 0.9998 ± 0.0002, respectively (Supplementary Figure 3H, Supplementary Table 1). These data further signify that the dropkick logistic regression model results in enhanced performance over one-dimensional heuristic thresholding, especially in the presence of high ambient noise in the training set.

### dropkick recovers expected cell populations and eliminates low-quality barcodes in experimental data

To evaluate dropkick’s performance against existing scRNA-seq filtering algorithms with real-world data, we processed a human T cell dataset from 10X Genomics (Figure 1) and again compared default dropkick results (dropkick_label) to 10X CellRanger version 2 and EmptyDrops (Lun, et al. 2019). The final dropkick coefficients and chosen regularization strength (lambda; Figure 4A) reveal that the model is sparse - with nearly 98 % of all coefficient values equal to zero – offering an interpretable gene-based output. Without prior training or supervision, dropkick identified higher counts of mitochondrial genes, which are markers of cell death and poor barcode quality (Tait and Green 2010), as predictive of empty droplets (Figure 4A). To visualize heuristic distributions within the T cell dataset, the number of genes detected and the percentage of ambient counts per barcode are shown along with dropkick’s automatic training thresholds (Figure 4B). Junk barcodes below the lower n_genes threshold were discarded before model training and assigned a dropkick score of zero. Barcodes between the two thresholds were initially assigned a label indicating putative empty droplets, while those above the upper threshold were labeled as real cells for model training. The dropkick score overlay illustrates how dropkick re-drew label boundaries in gene space (Figure 4B). dropkick scores are noticeably lower for barcodes with high ambient RNA content, while some putative empty droplets with lower background are “rescued” and labeled as real cells by the trained dropkick model.

**Figure 4.**
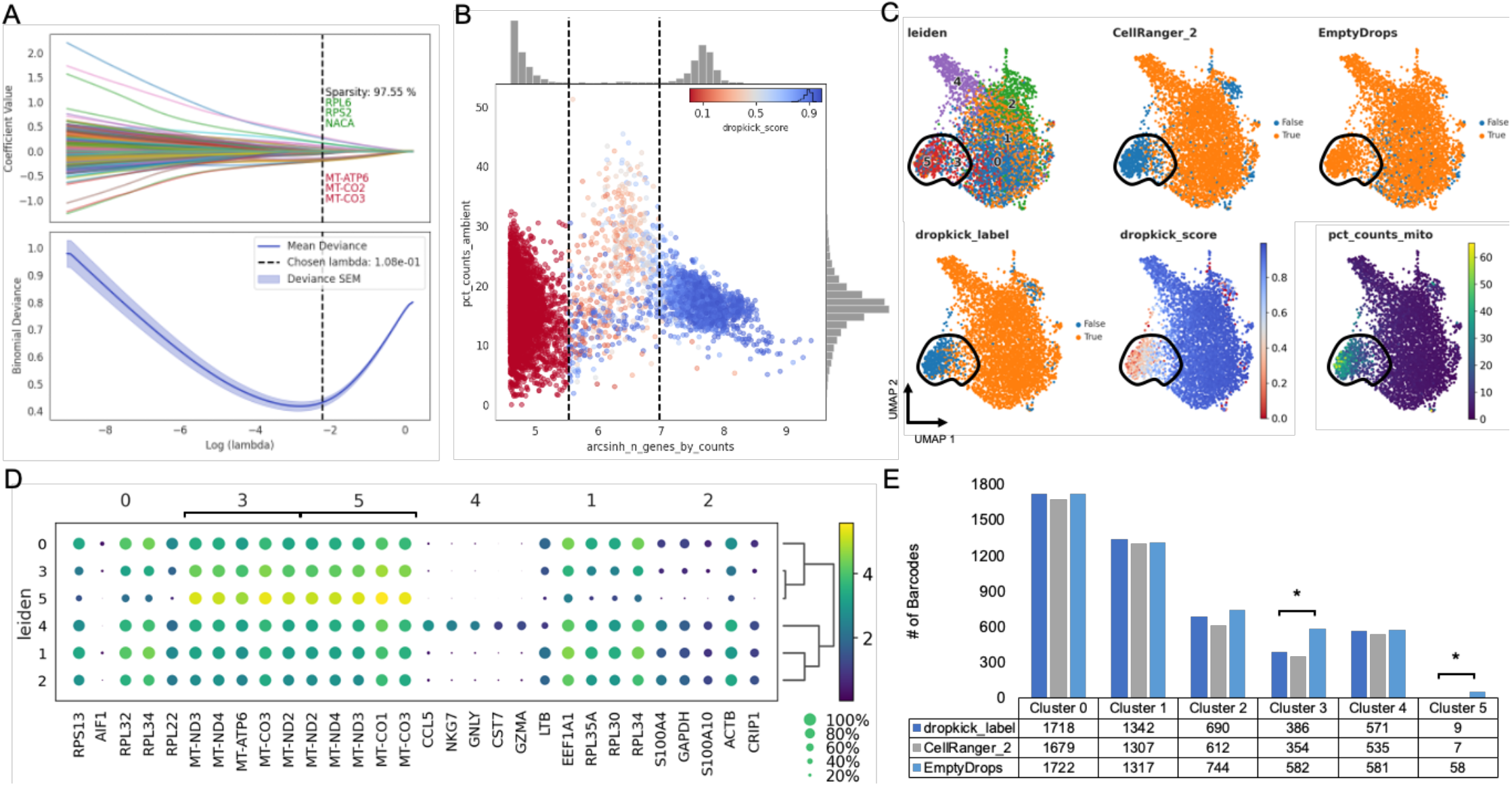
dropkick recovers expected cell populations and eliminates low-quality barcodes in experimental data. A) Plot of coefficient values for 2,000 highly variable genes (top) and mean binomial deviance ± SEM (bottom) for five-fold cross-validation along the lambda regularization path defined by dropkick. Top and bottom three coefficients are shown, in axis order, along with total model sparsity representing the percentage of coefficients with values of zero (top). Chosen lambda value indicated by dashed vertical line. B) Joint plot showing scatter of percent ambient counts versus arcsinh-transformed genes detected per barcode, with histogram distributions plotted on margins. Initial dropkick thresholds defining the training set are shown as dashed vertical lines. Each point (barcode) is colored by its final dropkick score after model fitting. C) UMAP embedding of all barcodes kept by dropkick_label (dropkick score ≥ 0.5), CellRanger_2 and EmptyDrops. Points colored by each of the three filtering labels, as well as clusters determined by NMF analysis, dropkick score (cell probability), and percent counts mitochondrial. Circled area shows high mitochondrial enrichment in a population discarded by dropkick. D) Dot plot showing top differentially expressed genes for each NMF cluster. The size of each dot indicates the percentage of cells in the population with nonzero expression for the given gene, while the color indicates the average expression value in that population. Bracketed genes indicate significantly enriched populations in EmptyDrops compared to dropkick_label as shown in E. E) Table and bar graph enumerating the total number of barcodes detected by each algorithm in all NMF clusters. EmptyDrops shows significant enrichment over dropkick_label in clusters 3 and 5 as determined by sc-UniFrac analysis.

We then jointly processed all barcodes kept by dropkick_label (dropkick score ≥ 0.5), CellRanger_2, and EmptyDrops using nonnegative matrix factorization (NMF; Kotliar, et al 2019) to define cell clusters, and sc-UniFrac (Liu, et al. 2018) to determine population differences across labeled barcode sets. A UMAP embedding of these barcodes reveals a population of cells with high mitochondrial content that is mostly excluded by dropkick (Figure 4C). This area is enriched in clusters 3 and 5 from NMF analysis, which carry exclusively mitochondrial genes as their top differentially expressed features (Figure 4D). Based on sc-UniFrac, these two clusters constitute the only statistically significant differences between EmptyDrops and dropkick (Figure 4E). Final dropkick labels had an overall sc-UniFrac distance of 0.01 from CellRanger_2 and 0.02 from EmptyDrops across all barcodes. These data indicate that dropkick recovers as many or more real cells in expected populations than previous algorithms, while also identifying and excluding low-quality dead or dying cells with high mitochondrial RNA content.

### dropkick outperforms analogous methods on challenging datasets

To challenge the robustness of the model, we next used dropkick to filter real-world samples with more complex cell types and higher noise. Human colorectal carcinoma (3907_S2) and adjacent normal colonic mucosa (3907_S1) samples were dissociated and encapsulated using the inDrop scRNA-seq platform (Klein, et al. 2015). In contrast to the 10X Genomics pan-T cell dataset (Figure 1, Figure 4), these samples exhibited high levels of background, containing empty droplets with thousands of UMIs detected per barcode and up to 40 % ambient RNA in expected cell barcodes (Supplementary Figure 5A,D). Because of this dominant ambient profile, infiltrating immune populations with lower mRNA content than epithelial cells can be lost among empty droplets. Indeed, CellRanger_2 and EmptyDrops show depletion in T cells (cluster 7) and macrophages (cluster 11) compared to dropkick (Figure 5A,B). Prevalence of high-RNA empty droplets also yields a population with low genetic diversity and mitochondrial gene enrichment (cluster 4; Figure 5A) that is kept by the one-dimensional thresholding of CellRanger_2 but discarded by dropkick. sc-UniFrac analysis confirmed that dropkick recovers significantly more cells from rare populations than both CellRanger_2 and EmptyDrops in this pair of high-background datasets dominated by ambient RNA from dead and dying colonic epithelial cells (Figure 5C, Supplementary Figure 5). Meanwhile, dropkick also identified and removed significantly more dead cells (cluster 4) than both CellRanger_2 and EmptyDrops (Figure 5C) by identifying mitochondrial and ambient genes as negative coefficients (Supplementary Figure 5B,E).

**Figure 5.**
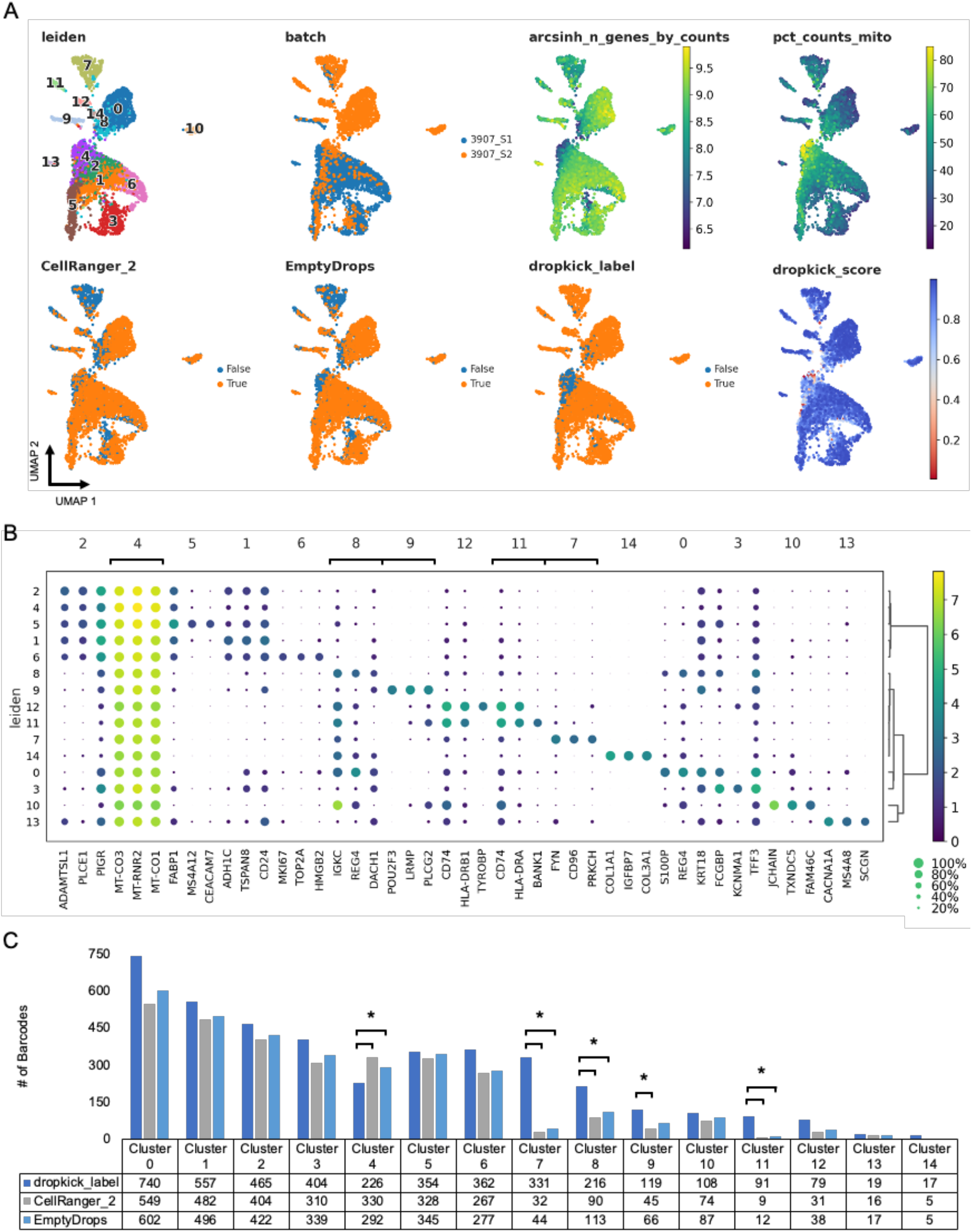
dropkick outperforms analogous methods on challenging datasets. A) UMAP embedding of all barcodes kept by dropkick_label (dropkick score ≥ 0.5), CellRanger_2 and EmptyDrops for human colorectal carcinoma inDrop samples. Points colored by each of the three filtering labels, as well as clusters determined by NMF analysis, dropkick score (cell probability), arcsinh-transformed total genes detected, percent counts mitochondrial, and original batch. 3907_S1 is normal human colonic mucosa and 3907_S2 is colorectal carcinoma from the same patient. B) Dot plot showing top differentially expressed genes for each NMF cluster. The size of each dot indicates the percentage of cells in the population with nonzero expression for the given gene, while the color indicates the average expression value in that population. Bracketed genes indicate significantly enriched or depleted populations in dropkick compared to CellRanger_2 and/or EmptyDrops labels as shown in C. C) Table and bar graph enumerating the total number of barcodes detected by each algorithm in all NMF clusters for the combined dataset. Significant cluster enrichment as determined by sc-UniFrac is denoted by brackets.

### dropkick filters reproducibly across scRNA-seq batches

We also applied dropkick to a combined human placenta dataset from six patients to show robustness of the model to batch-specific variation. dropkick learned the distribution of genes and ambient RNA specific to each dataset and filtered them accordingly (Supplementary Figure 6A), with a resulting AUROC of 0.9956 ± 0.0051 across all six replicates compared to EmptyDrops labels (Supplementary Figure 7E, Supplementary Table 3).

Extending this analysis to a larger cohort of scRNA-seq samples from both 10X Genomics (n = 13) and inDrop (n = 33) encapsulation platforms, we see that dropkick is highly concordant with CellRanger version 2 (AUROC 0.9656 ± 0.0271) and EmptyDrops (AUROC 0.9817 ± 0.012) results (Supplementary Figure 7A-D, Supplementary Table 3, Supplementary Table 4), suggesting global recovery of major cell populations. We also performed manual cell labelling to provide an additional alternative to compare to dropkick filtering on 33 inDrop datasets (see methods: CellRanger 2, EmptyDrops, and manual filtering of real-world scRNA-seq datasets). dropkick scores for these samples had an AUROC of 0.9729 ± 0.0335 compared to manually curated labels (Supplementary Figure 7E, Supplementary Table 5), again confirming the model’s utility for robust filtering across several unique datasets. Finally, we measured the total run time of dropkick for all of the above scRNA-seq batches, which was appreciably faster than the EmptyDrops R package, running to completion in 40.56 ± 25.97 seconds across ten replicates of all 46 samples when utilizing five CPUs with dropkick’s built-in parallelization (Supplementary Figure 7F).

## Discussion

Barcode filtering is a key preprocessing step in analyzing droplet-based single-cell expression data. Reliable filtering is confounded by distributions of global heuristics such as total counts, total genes, and ambient RNA that can be highly variable across batches and encapsulation platforms. We have developed dropkick, a fully automated machine learning software tool that assigns confidence scores and labels to barcodes from unfiltered scRNA-seq counts matrices. By automatically curating a training set using predictive heuristics and training a gene-based logistic regression model, dropkick ensures that ambient barcodes (“empty droplets”) are removed from the filtered dataset while recovering rare, low-RNA cell types that may be lost in ambient noise. We showed that unlike previous filtering approaches including one-dimensional thresholding (CellRanger 2) and a Dirichlet-multinomial model (EmptyDrops), dropkick is robust to the level of ambient RNA, performing favorably in both low and high-background scenarios across simulated and real-world datasets.

Although we have demonstrated that dropkick is more robust to varying degrees of ambient background than existing filtering methods, the dropkick model is still limited by the input dataset. As stated previously (see Evaluating dataset quality with dropkick QC module), the profile of ranked total counts/genes and the global contribution of ambient reads are vital to analysis of single-cell sequencing data, including cell filtering. Data with weak separation between high-quality cells and empty droplets (i.e. a unimodal distribution of n_genes lacking distinct plateaus in the log-rank curve) will perform poorly in inflection-point thresholding as well as data-driven models such as EmptyDrops and dropkick due to the similarity between theoretically “high-confidence” barcodes and ambient background droplets. Moreover, datasets dominated by a small number of ambient genes (> 40 % average ambient counts across all barcodes) will also perform poorly in automated filtering. While such data artefacts may be handled by dropkick’s heavy feature selection conferred by HVG calculation and elastic net regularization, there will also be circumstances that cause dropkick – as well as CellRanger and EmptyDrops - to return an over or under-filtered dataset. Scenarios such as those described should be considered QC failures, and further analysis should not be performed. For this reason, the dropkick QC module is extremely beneficial in post-alignment evaluation of scRNA-seq data quality and should be applied to all datasets prior to filtering. The dropkick Python package provides a fast, user-friendly interface that integrates seamlessly with the Scanpy (Wolf, et al. 2018) single-cell analysis suite for ease of workflow implementation. dropkick is available for install through the Python Package Index (pypi.org/project/dropkick/), and source code is hosted on GitHub (github.com/KenLauLab/dropkick).

## Methods

### Quality control and ambient RNA quantification with the dropkick QC module

The dropkick QC module begins by calculating global heuristics per barcode (observation) and gene (variable) using the scanpy (Wolf, Angerer, and Theis 2018) *pp.calculate_qc_metrics* function. These metrics are used to order barcodes by decreasing total counts (black curve in Figure 1A) and order genes by increasing dropout rate (Figure 1B). The *n*th gene ranked by dropout rate determines the cutoff for calling “ambient” genes, with *n* determined by the n_ambient parameter in the *dropkick.qc_summary* function. All genes with dropout rates less than or equal to this threshold are labeled “ambient”. In a sample with many (> *n*) genes detected in all barcodes, this ensures that the entire ambient profile is identified. Through observation of samples used in this study, we set the default n_ambient = 10. To compile the dropkick QC summary report, the log-total counts versus log-ranked barcodes (Figure 1A black curve) are plotted along with total genes detected for each barcode (Figure 1A green points), percent counts from “ambient” genes in each barcode (Figure 1A blue points), and percent counts from mitochondrial genes in each barcode (Figure 1A red points).

### Labeling training set with the dropkick filtering module

The dropkick filtering module also begins by calculating global heuristics per barcode (observation) and gene (variable) using the scanpy (Wolf, Angerer, and Theis 2018) *pp.calculate_qc_metrics* function. Next, training thresholds are calculated on the histogram of the chosen heuristic(s); arcsinh-transformed n_genes by default. dropkick then uses the scikit-image function *filters.threshold_multiotsu* to identify two local minima in the n_genes histogram that represent the transitions from “junk” barcodes to “empty droplets” and from “empty droplets” to real cells. These locations are also characterized by the two expected drop-offs in the total counts/genes profiles as shown in the dropkick QC report (Figure 1, Supplementary Figure 1). To label barcodes for dropkick model training, barcodes with fewer genes detected than the first multi-Otsu threshold are discarded due to their lack of molecular information. dropkick then labels barcodes below the second threshold as “empty”, and remaining barcodes above the second threshold as real cells for initial training. These inputs to the dropkick logistic regression model represent the “noisy” boundary in heuristic space that is to be replaced with a learned cell boundary in gene space.

### Training and optimizing the dropkick filtering model

The dropkick filtering model uses logistic regression with elastic net regularization (Zou and Hastie 2005), and is fit as described in Friedman, et al. 2010. The elastic net combines ridge and lasso (least absolute shrinkage and selection operator) penalties for optimal regularization of model coefficients. The ridge regression penalty pushes all coefficients toward zero while allowing multiple correlated predictors to borrow strength from one another, ideal for a scenario like scRNA-seq with several expected collinearities (Hoerl and Kennard 1970). The lasso penalty on the other hand, favors model sparsity, driving coefficients to zero and thus selecting informative features (Tibshirani 1996). The combined elastic net balances feature selection and grouping by preserving or removing correlated features from the model in concert (Zou and Hastie 2005).

The fraction *α* ∈ [0,1] (alpha) represents the balance between the lasso and ridge penalties. If *α* = 0, the regularization would be entirely ridge, while if a = 1, it would be entirely lasso. By default, dropkick fixes this alpha value at 0.1, but the user may alter this parameter or provide multiple alpha values to optimize through cross-validation (with lambda; explained below) at the expense of slightly longer computational time.

For a desired length of “lambda path,” *n* (default *n* = 100 for dropkick), the model is fit *n* + 1 times, where the first pass determines the values of lambda (regularization strength) to test, and subsequent fits determine model performance using cross-validation (CV; default 5-fold for dropkick). Each fit involves selection of highly-variable genes (HVGs; scanpy *pp.highly_variable_genes;* default 2,000 for dropkick) from the training set. For both the first pass and the final model, the training set consists of all available barcodes, while training the model along the lambda path uses only the current training fold as to not bias model fitting with information from the test set. The lambda path is scored using mean deviance from the training labels for all cross-validation folds. The largest value of lambda such that its mean CV deviance is less than or equal to one standard error above the minimum deviance is chosen as the final regularization strength for the model in order to further minimize overfitting. Finally, dropkick fits a logistic regression model using all training labels and the chosen lambda value and assigns cell probability (dropkick_score) to all barcodes. By default, the resulting dropkick_label is positive (1; real cell) for barcodes with dropkick_score ≥ 0.5, but the user may define a stricter or more lenient threshold for particular applications.

### scRNA-seq data simulation

We used CellBender (Fleming, et al. 2019) to simulate single-cell datasets. We generated a basic count matrix with 30,000 features (n_genes), 12,000 total droplets (including 3,000 n_cells and 9,000 n_empty), and 6 clusters. The default ratio between the cell size scale factor and the empty droplet size scale factor – d_cell at 10,000 and d_empty at 200 – created an unrealistic gap between the empty droplets and the real cells but built a foundation on which to produce more realistic simulations. By adjusting these parameters, we simulated two different scenarios with the number of features, total droplets, and clusters held constant. The first scenario modeled a “low background” dataset, with a realistic n_genes and total counts profile and relatively low ambient RNA. We set the cell size scale factor (d_cell) to 10,000, and the empty droplet size scale factor (d_empty) to 1,000. These settings produced a small gap between the real cells and the empty droplets, yet still mimicked a low background droplet profile. We then modeled a “high background” scenario, which had much higher ambient RNA content. For this simulation we set d_cell to 10,000, and d_empty to 2,000. This simulation mimicked a real scRNA-seq dataset with a high ambient profile, as it had a smaller gap between real cells and empty droplets. Taken together, these simulations recapitulate real-world single-cell data and were tested by dropkick to compare their ground-truth labels to those determined by dropkick filtering.

### CellRanger 2, EmptyDrops, and manual filtering of real-world scRNA-seq datasets

CellRanger filtering algorithms were derived from Lun, et al. 2019, with CellRanger 2 described by the function *DefaultDrops* (from the repository github.com/MarioniLab/EmptyDrops2017), and EmptyDrops by the *EmptyDrops* function within the DropletUtils R package (v1.8.0). All 10X datasets, with the exception of 3659_colon, were processed as in Lun, et al. 2019 (github.com/MarioniLab/EmptyDrops2017). EmptyDrops was run for the 3659_colon sample not present in Lun et al. 2019 using a minimum non-ambient counts threshold of 500 UMIs. For inDrop datasets, CellRanger 2 was run with an expected cell number of 1,000, and upper_quant and lower_prop parameters set to 0.99 and 0.01, respectively. EmptyDrops was run for all inDrop datasets using the “inflection point” as the minimum non-ambient counts threshold as in Lun, et al. 2019 (github.com/MarioniLab/EmptyDrops2017). Manual filtering was performed for each inDrop sample by initial thresholding beyond the “knee point” detected in the first curve of the ranked barcodes profile (as in Figure 1A). Then, following standard dimension reduction and high-resolution Leiden clustering, clusters with low quality cells (high mitochondrial/ambient percentage, low total counts/genes) were manually gated out of the final dataset. These manually curated labels were used as an orthogonal “gold standard” for benchmarking automated thresholding methods (Supplementary Figure 2A) and final AUROC (Supplementary Figure 4D).

### sc-UniFrac analysis of shared populations between dropkick, CellRanger 2, and EmptyDrops labels

In order to evaluate the preservation of expected cell clusters between dropkick and alternative labels, we employed sc-UniFrac (Liu, et al. 2018) to determine the global and populational differences between the label sets. We used nonnegative matrix factorization (NMF) to analyze the union of barcodes kept by dropkick_label, CellRanger_2, and EmptyDrops in order to reduce dimensions into cell identity and activity “metagenes” (Kotliar, et al. 2019). We then clustered this low-dimensional space using the Leiden algorithm (Traag, et al. 2019) to define consensus cell populations for sc-UniFrac analysis. We then ran sc-UniFrac (v0.9.6) to evaluate statistically significant cluster differences based on both cluster membership and gene expression hierarchies between clusters. The global sc-UniFrac distance quantified the overall similarity of hierarchical trees across barcode label sets.

### Dimension reduction, clustering, projection, and differential expression analysis

We used Consensus Nonnegative Matrix Factorization (cNMF; Kotliar, et al. 2019) for initial dimension reduction. The optimal number of factors, *k*, was determined by maximizing stability and minimizing error across all tested values after 30 iterations of each. We then built a nearest-neighbors graph in Scanpy (*pp.neighbors* function) from the NMF usage scores for consensus factors in all cells, where we set n_neighbors to the square root of the total number of cells in the dataset. We then clustered cells with the Leiden algorithm (Scanpy *tl.leiden* function; Traag, Waltman, and Van Eck 2019) applied to this graph. Resulting clusters were used in sc-UniFrac analysis, differential expression, and visualization. We performed differential expression analysis using a Student’s t-test with Benjamini-Hochberg p-value correction for multiple testing (Scanpy *tl.rank_genes_groups*). To visualize datasets in 2D space, we ran partition-based graph abstraction (PAGA; Wolf, et al. 2019; Scanpy *tl.paga*) on this nearest-neighbors graph and associated Leiden clustering in order to create a simple representation of cluster similarity. Finally, a UMAP projection (McInnes and Healy 2018) seeded with these PAGA positions provided a two-dimensional embedding of all cells in the dataset (Scanpy *tl.umap* with init_pos=“paga”).

## Data and Code Availability

All publicly available datasets are listed in Supplementary Table 6, including the human colorectal carcinoma inDrop data deposited to the Gene Expression Omnibus (GEO) to accompany this manuscript (GSE158636). inDrop scRNA-seq data were generated with according to published protocols (Southard-Smith, et al. 2020; Banerjee et al., 2020). The dropkick Python package is available for download via “pip” from the Python Package Index (PyPI) at https://pypi.org/project/dropkick/. Source code for the package is also available on GitHub at https://github.com/KenLauLab/dropkick. Scripts for reproducing analyses in this manuscript are hosted on GitHub at https://github.com/codyheiser/dropkick-manuscript.

## Acknowledgments

The authors would like to acknowledge the Epithelial Biology Center and Vanderbilt Quantitative Systems Biology Center for helpful discussions, and Paige Vega from the Lau Lab for assistance in tailoring the sc-UniFrac analysis pipeline. K.S.L. is funded by the NIH (grants R01DK103831, P50CA236733, U01CA215798, and U54CA217450), and C.N.H. is funded by NIH grant U2CCA233291.

## Author Contributions

B.C. and C.N.H. conceived of the quality control and filtering methodology. J.J.H. assisted in design and interpretation of the statistical model and analysis. C.N.H. developed the dropkick software package and analyzed the data. V.M.W. performed simulations and sc-UniFrac analyses. C.N.H. and V.M.W. wrote the manuscript. K.S.L. supervised the study, secured funding, and participated in writing the manuscript and interpreting results.

## Declaration of Interests

The authors declare no competing interests.

## Supplementary Figures, Tables and Legends

**Supplementary Figure 1.**
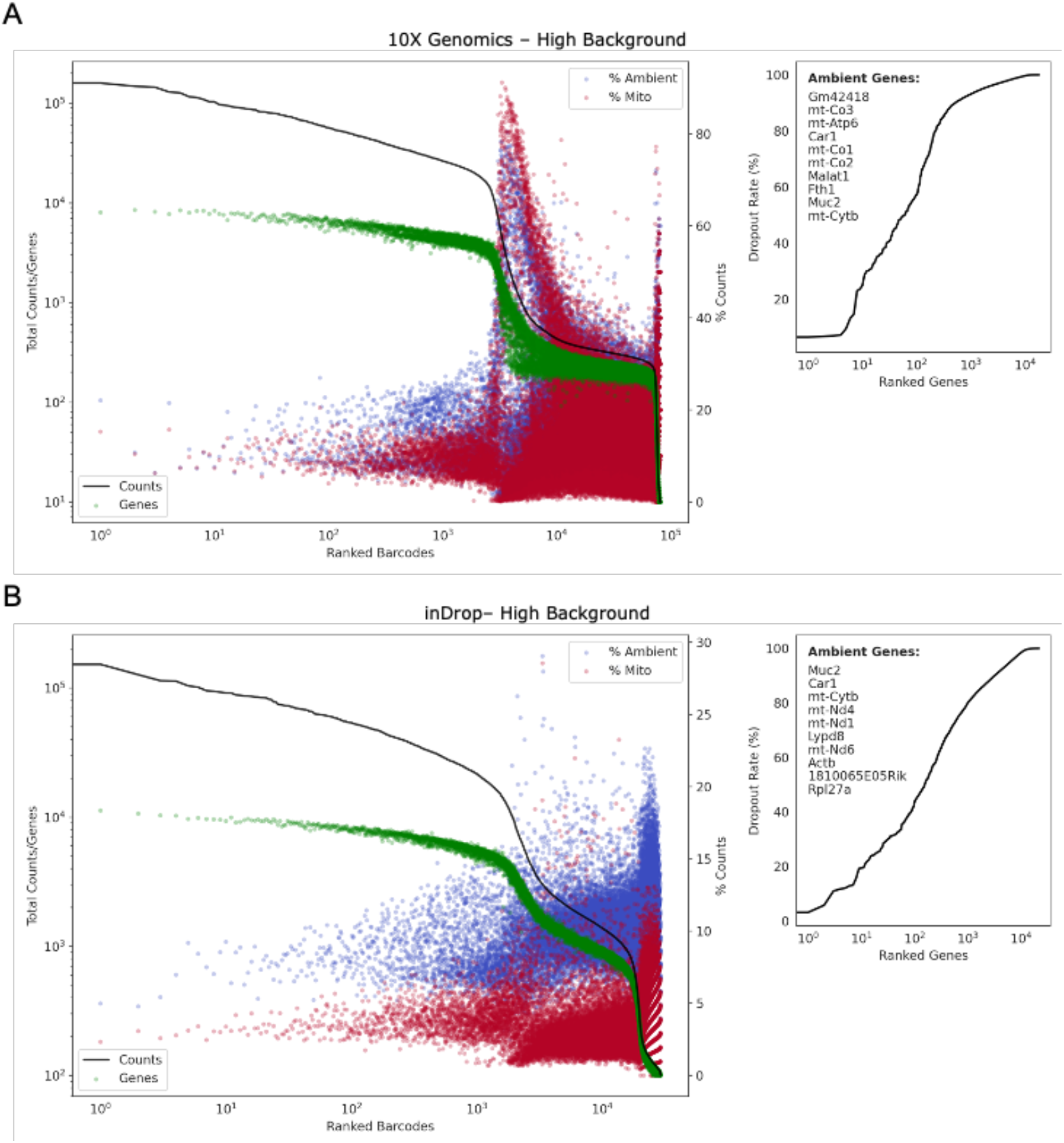
dropkick QC reports for a mouse colonic epithelium sample analyzed by both 10X Genomics (A) and inDrop (B) scRNA-seq. In contrast to Figure 1, this is considered a high-background sample due to the height (increased total counts) of the second plateau (empty droplets) and presence of epithelial marker genes (*Car1, Muc2*) in the ambient profile.

**Supplementary Figure 2.**
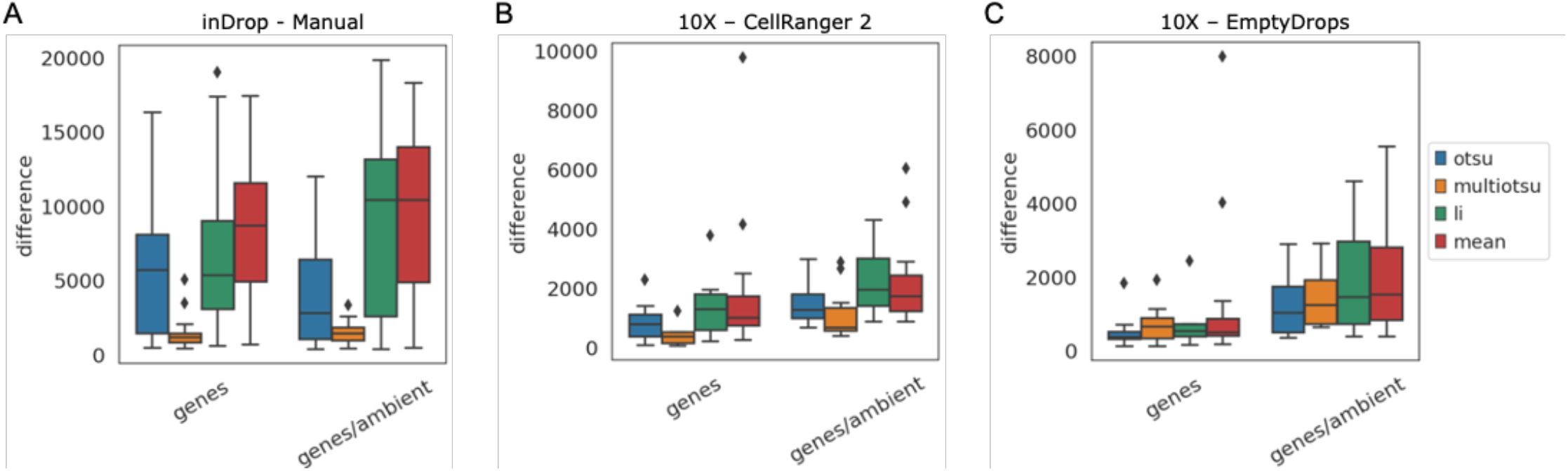
Optimal heuristics and thresholding for determination of dropkick training set. A) Barcode set differences between initial dropkick thresholding and manual filtering of 33 inDrop scRNA-seq datasets. Four automated thresholding techniques were used to label cells based on the distribution of arcsinh-transformed genes detected alone (genes), or the combination of genes and percent ambient counts as calculated by the dropkick QC module (genes/ambient; see Methods: Quality control and ambient RNA quantification with dropkick QC module). B) Same as in A for 13 10X Genomics scRNA-seq datasets, with set differences compared to CellRanger_2. C) Same as in B, with set differences compared to EmptyDrops.

**Supplementary Figure 3.**
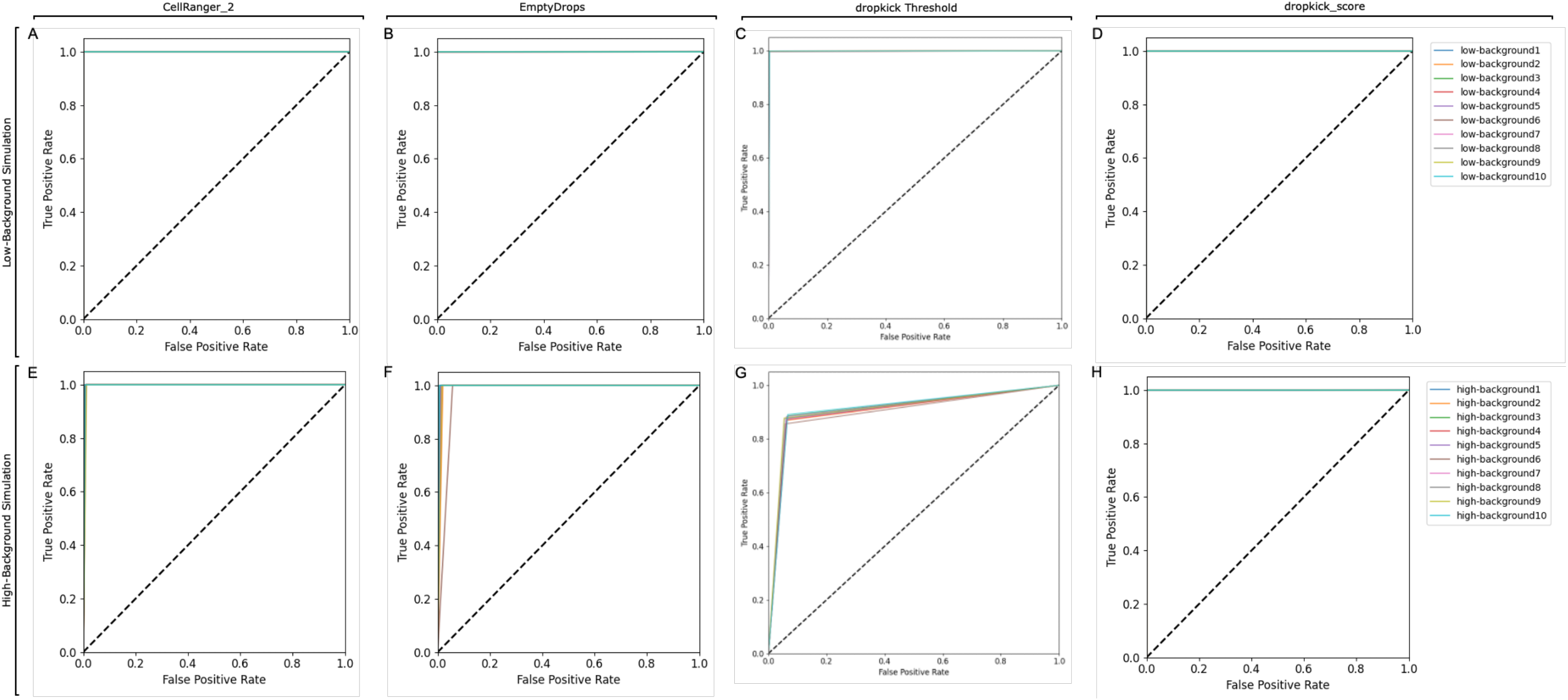
Testing dropkick on simulated datasets. A) Receiver operating characteristic (ROC) curves for CellRanger_2 vs. ground truth in ten low-background simulations. B) ROC curves for EmptyDrops vs. ground truth in ten low-background simulations. C) ROC curves for dropkick training labels (threshold) vs. ground truth in ten low-background simulations. D) ROC curves for final dropkick score vs. ground truth in ten low-background simulations. E-H) Same as in A-D, for ten high-background simulations.

**Supplementary Figure 4.**
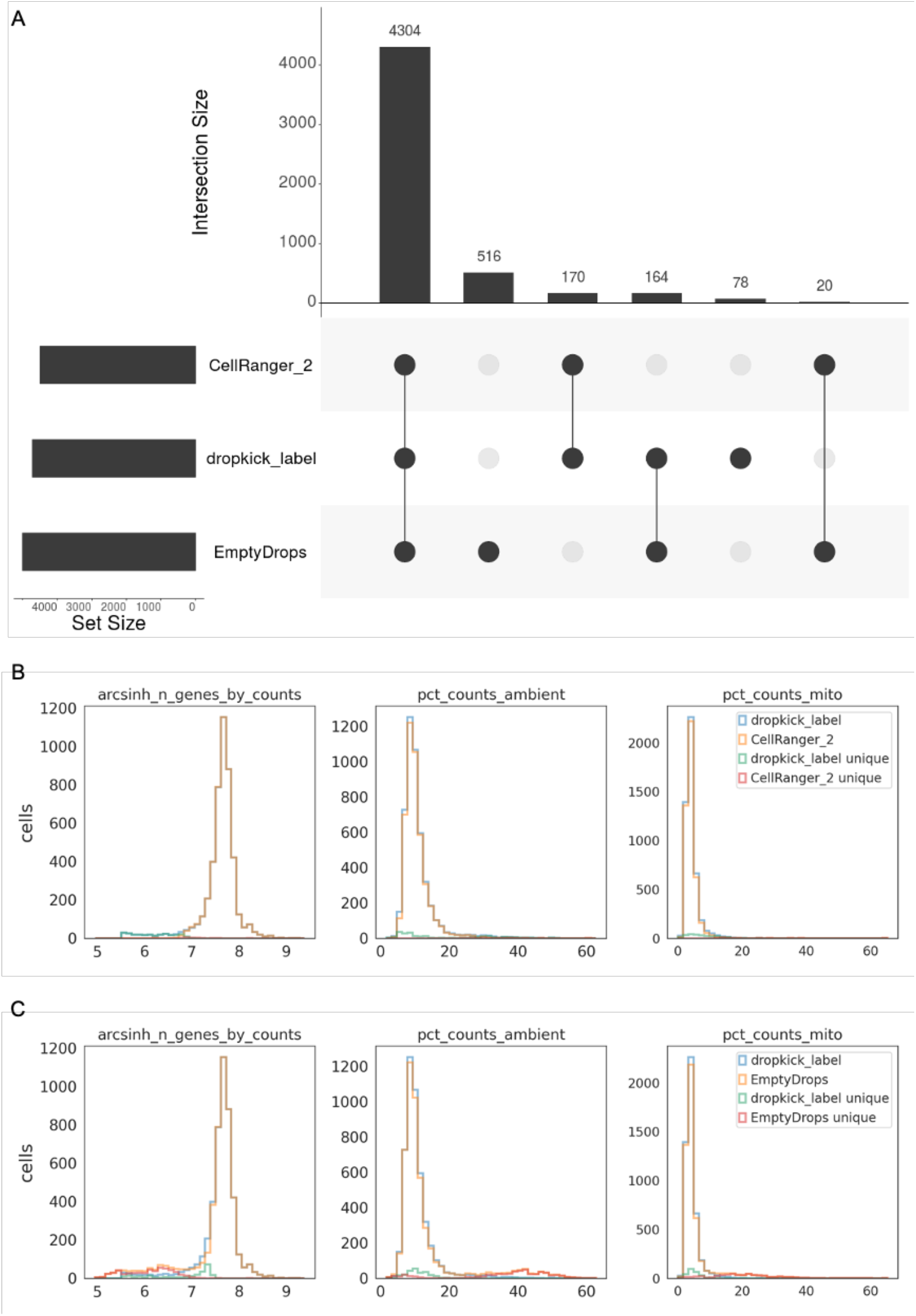
Barcode set differences for 4k pan-T cell dataset. A) UpSet plot showing global set differences between dropkick_label (dropkick score ≥ 0.5), CellRanger_2 and EmptyDrops. B) Histograms showing global distribution of heuristics (arcsinh-transformed genes, left, percent ambient counts, middle, and percent mitochondrial counts, right) in barcodes kept by dropkick_label and CellRanger_2. Distribution of barcodes unique to each label set also overlaid to show difference. C) Same as in B, for dropkick_label compared to EmptyDrops.

**Supplementary Figure 5.**
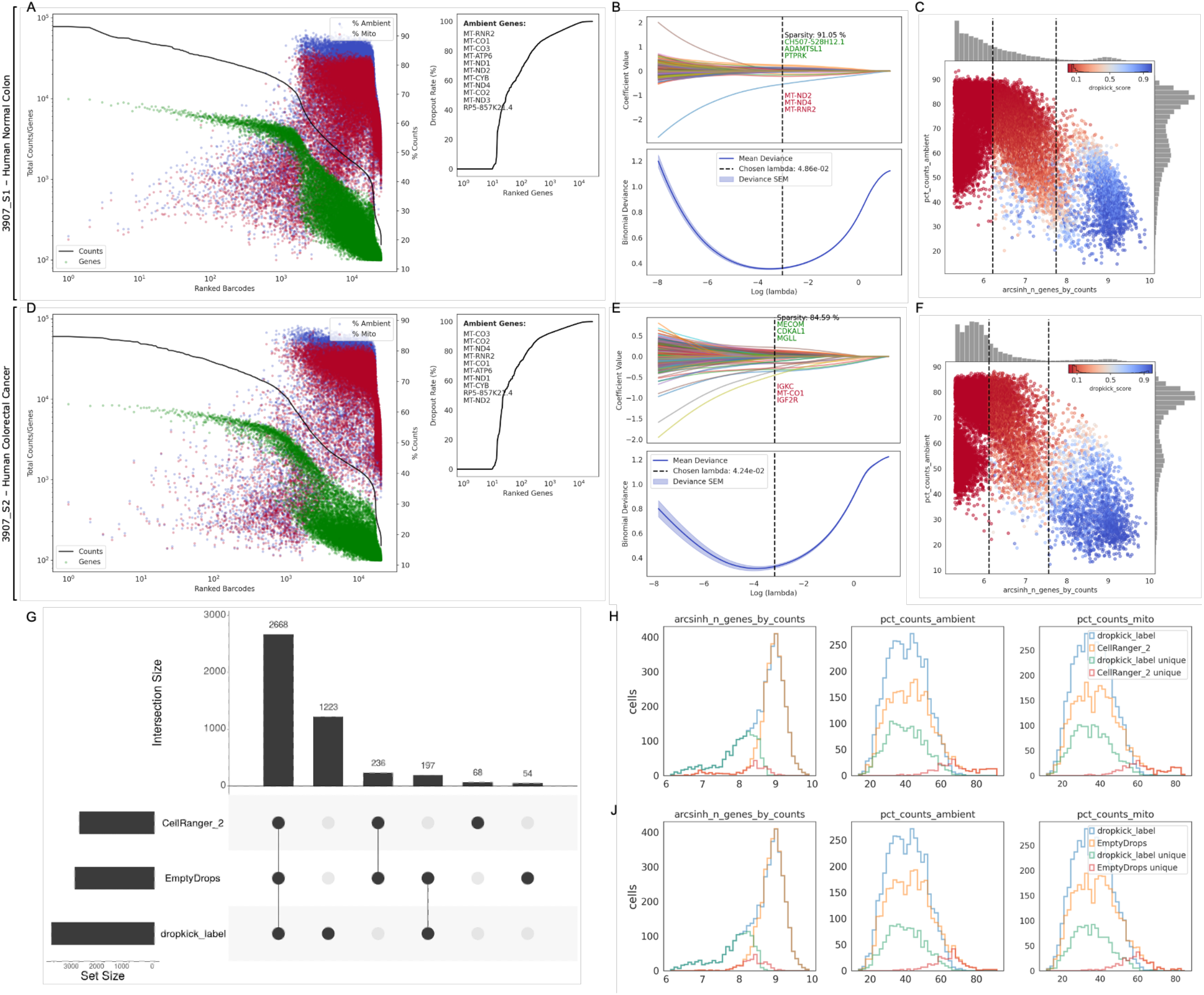
dropkick plots and barcode set differences for human colorectal carcinoma (CRC) inDrop samples. A) dropkick QC report for human normal colonic mucosa, 3907_S1 and CRC, 3907_S2. B) dropkick coefficient plots, showing coefficient values (top) and binomial deviance (bottom) along the tested lambda regularization path. Dashed line indicates chosen lambda value of trained model. Top and bottom three genes by coefficient value and total model sparsity noted in top plot. C) dropkick score plot showing scatter of percent counts ambient versus arcsinh-transformed total genes detected per barcode. Dashed lines indicate location of automated dropkick thresholds used for model training. Points colored by final dropkick score. D-F) Same as in A-C, but for adjacent human normal colonic mucosa sample, 3907_S2. G) UpSet plot showing global set differences between dropkick_label (dropkick score ≥ 0.5), CellRanger_2 and EmptyDrops. H) Histograms showing global distribution of heuristics (arcsinh-transformed genes, left, percent ambient counts, middle, and percent mitochondrial counts, right) in barcodes kept by dropkick_label and CellRanger_2. Distribution of barcodes unique to each label set also overlaid to show difference. J) Same as in H, for dropkick_label compared to EmptyDrops.

**Supplementary Figure 6.**
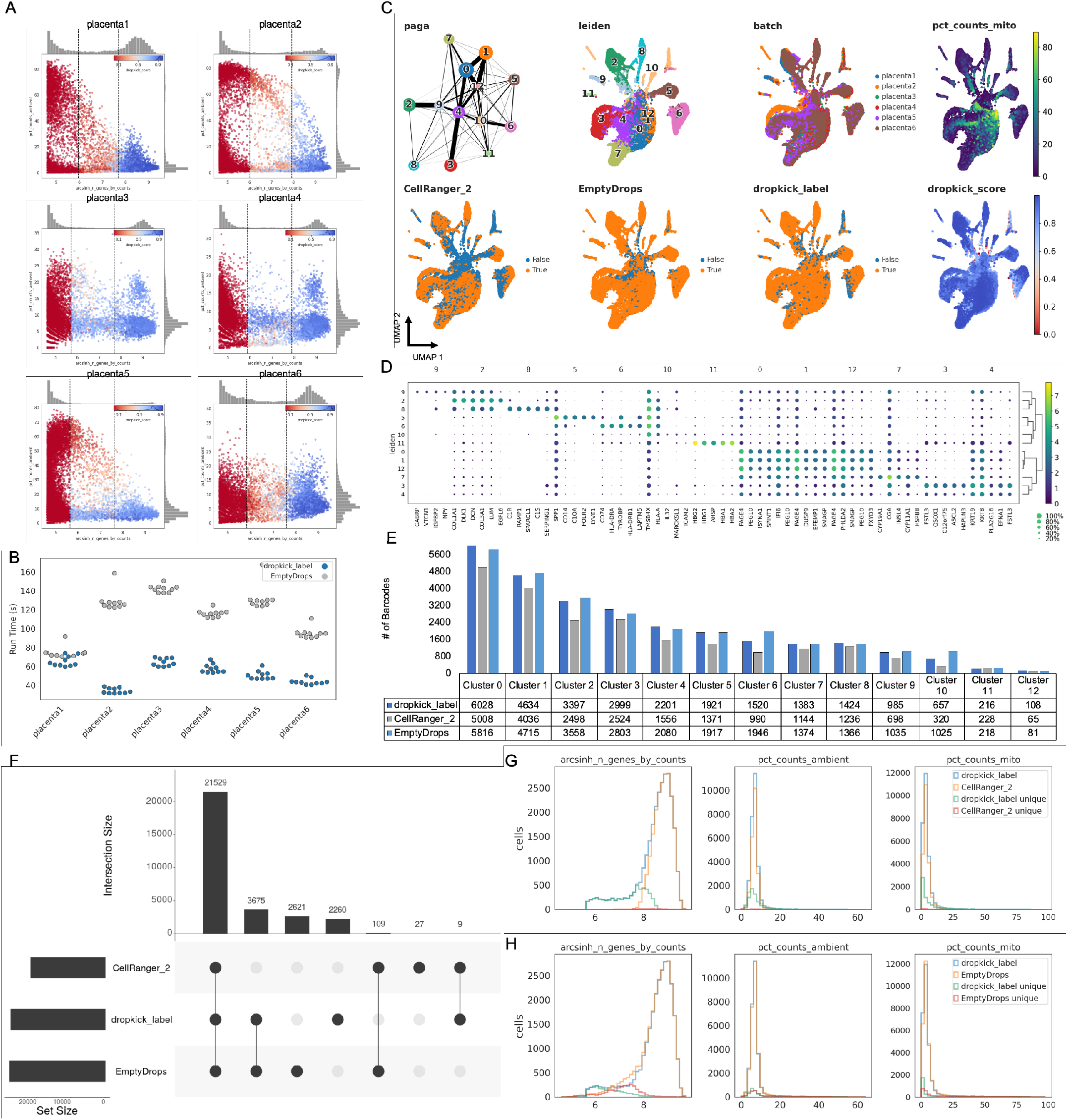
dropkick filters reproducibly across scRNA-seq batches. A) dropkick score plots for six placenta replicates. B) Total run time, in seconds, for EmptyDrops and dropkick. Both algorithms were run ten times on each placenta replicate; points represent single runs. C) PAGA graph and UMAP embedding of all barcodes kept by dropkick_label (dropkick score ≥ 0.5), CellRanger_2 and EmptyDrops for the aggregate placenta dataset. Points colored by each of the three filtering labels as well as original batch, NMF clusters, dropkick_score (cell probability), and percent counts mitochondrial. D) Dot plot showing top five differentially expressed genes for each NMF cluster. The size of each dot indicates the percentage of cells in the population with nonzero expression for the given gene, while the color indicates the average expression value in that population. E) Table and bar graph enumerating the total number of barcodes detected by each algorithm in all NMF clusters. F) UpSet plot showing global set differences between dropkick_label, CellRanger_2 and EmptyDrops. G) Histograms showing global distribution of heuristics (arcsinh-transformed genes, left, percent ambient counts, middle, and percent mitochondrial counts, right) in barcodes kept by dropkick_label and CellRanger_2. Distribution of barcodes unique to each label set also overlaid to show difference. H) Same as in G, for dropkick_label compared to EmptyDrops.

**Supplementary Figure 7.**
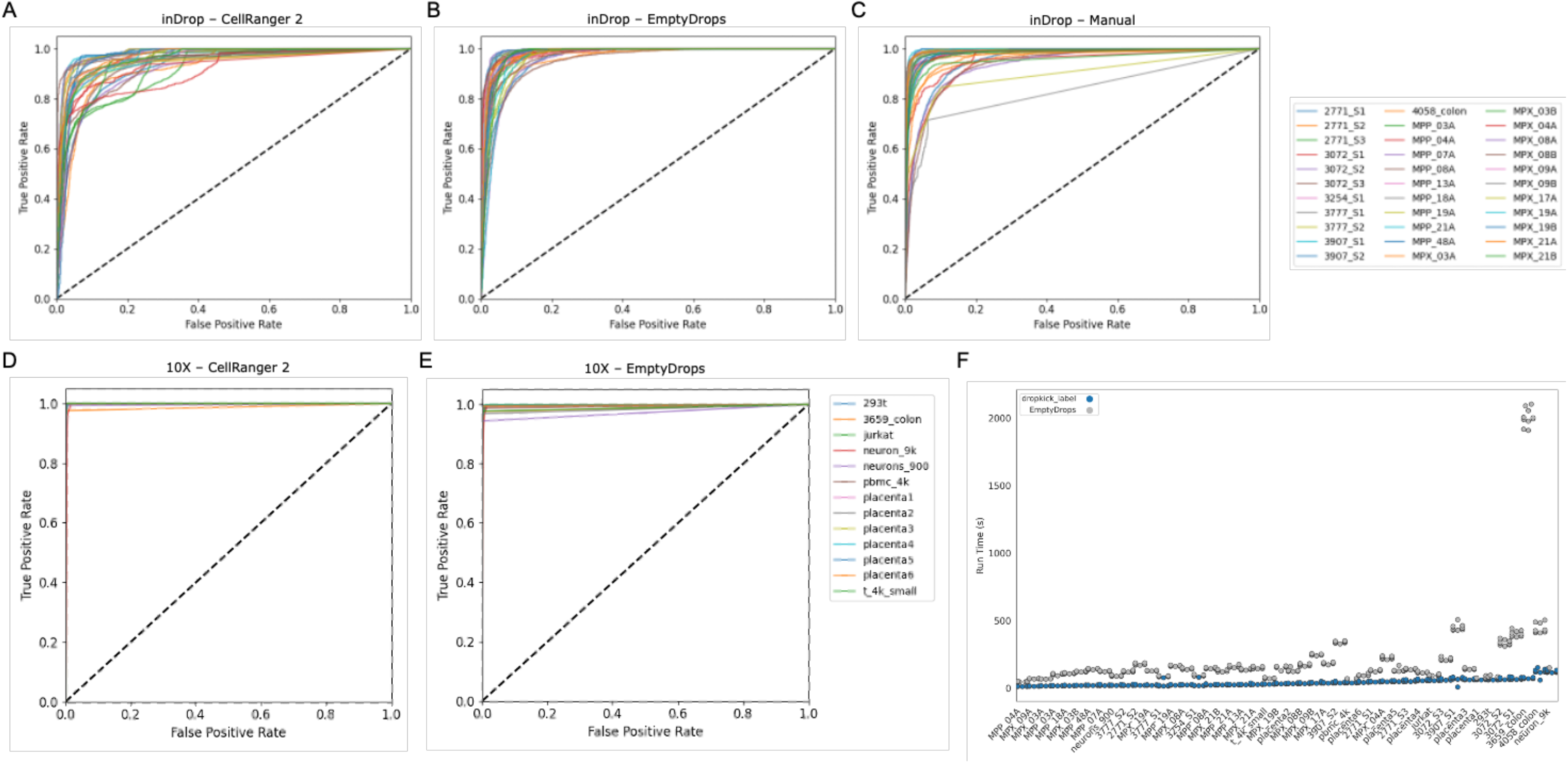
Comparing dropkick probability scores to two alternative cell labels using receiver operating characteristic (ROC) curves. AUC = area under the ROC curve. A) ROC curves for 13 10X Genomics scRNA-seq datasets, using CellRanger_2 as reference. B) Same as in A, with EmptyDrops as reference. C) ROC curves for 33 inDrop scRNA-seq datasets, using CellRanger_2 as reference. D) Same as in C, with EmptyDrops labels as reference. E) Same as in C, with manually curated cell labels as reference (see methods: CellRanger 2, EmptyDrops, and manual filtering of real-world scRNA-seq datasets). F) Total run time, in seconds, for EmptyDrops and dropkick. Both algorithms were run ten times on all datasets; points represent single replicates.

**Supplementary Table 1.** Global comparison statistics between automated thresholding (dropkick training labels) and trained dropkick model vs. ground-truth cell labels for low and high-background simulations.

| Simulation | Rep. | Threshold total cells | dropkick label total cells | True label total cells | Threshold sensitivity | Threshold Specificity | Threshold AUC | dropkick Sensitivity | dropkick Specificity | dropkick AUC |
| --- | --- | --- | --- | --- | --- | --- | --- | --- | --- | --- |
| Low Background | 1 | 3028 | 3001 | 3000 | 0.9983 | 0.9963 | 0.9973 | 1.0000 | 0.9999 | 1.0000 |
|  | 2 | 3022 | 3003 | 3000 | 0.9990 | 0.9972 | 0.9981 | 1.0000 | 0.9997 | 1.0000 |
|  | 3 | 3017 | 3003 | 3000 | 0.9990 | 0.9978 | 0.9984 | 1.0000 | 0.9997 | 1.0000 |
|  | 4 | 3022 | 3002 | 3000 | 0.9967 | 0.9965 | 0.9966 | 1.0000 | 0.9998 | 1.0000 |
|  | 5 | 3026 | 3001 | 3000 | 0.9990 | 0.9968 | 0.9979 | 1.0000 | 0.9999 | 1.0000 |
|  | 6 | 3023 | 3005 | 3000 | 0.9987 | 0.9970 | 0.9978 | 1.0000 | 0.9994 | 1.0000 |
|  | 7 | 3028 | 3003 | 3000 | 0.9983 | 0.9963 | 0.9973 | 1.0000 | 0.9997 | 1.0000 |
|  | 8 | 3012 | 3000 | 3000 | 0.9983 | 0.9981 | 0.9982 | 1.0000 | 1.0000 | 1.0000 |
|  | 9 | 3020 | 3000 | 3000 | 0.9990 | 0.9975 | 0.9982 | 1.0000 | 1.0000 | 1.0000 |
|  | 10 | 3030 | 3002 | 3000 | 0.9993 | 0.9965 | 0.9979 | 1.0000 | 0.9998 | 1.0000 |
| High Background | 1 | 3269 | 3005 | 3000 | 0.8910 | 0.9379 | 0.9124 | 1.0000 | 0.9994 | 1.0000 |
|  | 2 | 3178 | 2998 | 3000 | 0.8693 | 0.9404 | 0.9030 | 0.9990 | 0.9999 | 0.9995 |
|  | 3 | 3246 | 3002 | 3000 | 0.8860 | 0.9387 | 0.9103 | 1.0000 | 0.9998 | 1.0000 |
|  | 4 | 3167 | 3001 | 3000 | 0.8710 | 0.9420 | 0.9047 | 0.9997 | 0.9998 | 0.9998 |
|  | 5 | 3206 | 3002 | 3000 | 0.8833 | 0.9418 | 0.9108 | 1.0000 | 0.9998 | 1.0000 |
|  | 6 | 3095 | 2999 | 3000 | 0.8563 | 0.9448 | 0.8989 | 0.9997 | 1.0000 | 0.9998 |
|  | 7 | 3200 | 2999 | 3000 | 0.8737 | 0.9396 | 0.9047 | 0.9993 | 0.9999 | 0.9997 |
|  | 8 | 3127 | 2998 | 3000 | 0.8760 | 0.9475 | 0.9103 | 0.9993 | 1.0000 | 0.9997 |
|  | 9 | 3118 | 2998 | 3000 | 0.8780 | 0.9490 | 0.9121 | 0.9993 | 1.0000 | 0.9997 |
|  | 10 | 3199 | 2998 | 3000 | 0.8773 | 0.9407 | 0.9072 | 0.9990 | 0.9999 | 0.9995 |

**Supplementary Table 2.**
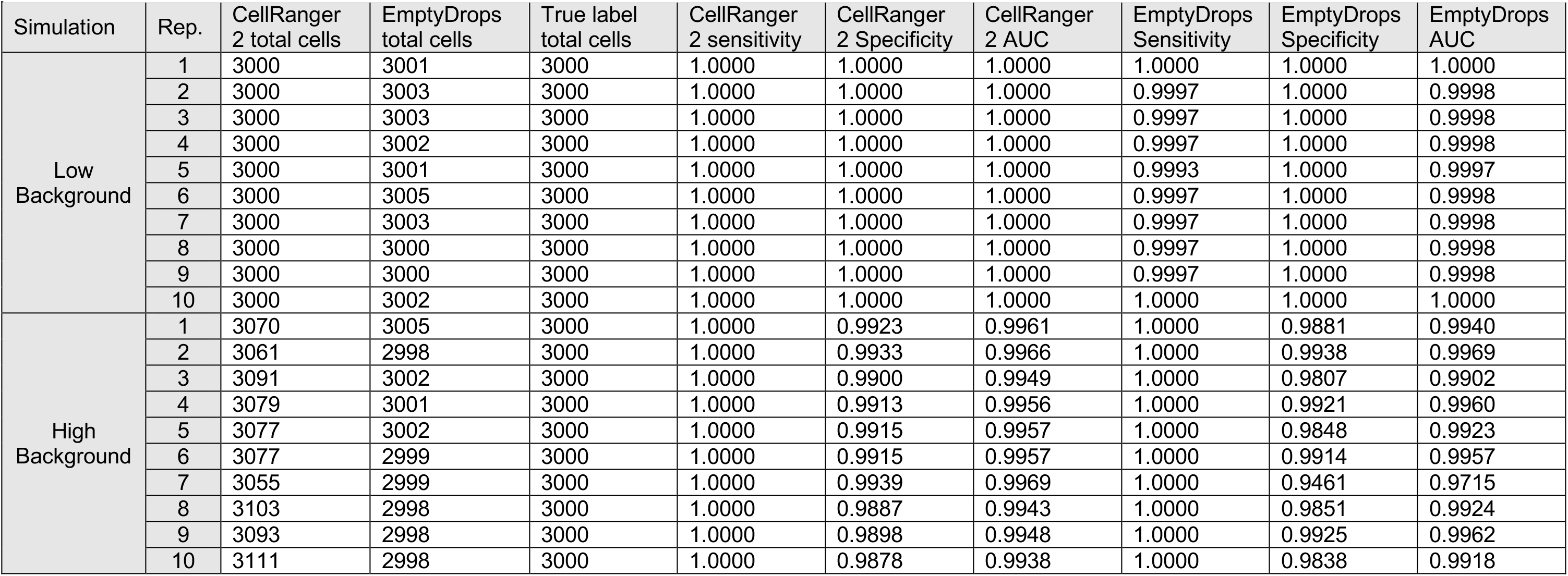
Global comparison statistics between CellRanger_2 and EmptyDrops versus ground-truth cell labels for low and high-background simulations.

**Supplementary Table 3.** Global comparison statistics between dropkick, CellRanger_2, and EmptyDrops for 13 10X Genomics scRNA-seq datasets.

|  | dropkick label total cells | CellRanger 2 total cells | CellRanger 2 sensitivity | CellRanger 2 specificity | CellRanger 2 AUC | EmptyDrops total cells | EmptyDrops sensitivity | EmptyDrops specificity | EmptyDrops AUC |
| --- | --- | --- | --- | --- | --- | --- | --- | --- | --- |
| 293t | 3224 | 2897 | 0.9717 | 0.9986 | 0.9994 | 3028 | 0.9508 | 0.9988 | 0.9991 |
| jurkat | 4020 | 3259 | 0.9963 | 0.9974 | 0.9989 | 3452 | 0.9655 | 0.9977 | 0.9985 |
| neuron_9k | 11050 | 9094 | 0.8745 | 0.9958 | 0.9964 | 11027 | 0.8422 | 0.9976 | 0.9950 |
| neurons_900 | 1297 | 792 | 0.9369 | 0.9992 | 0.9963 | 1940 | 0.6649 | 1.0000 | 0.9721 |
| pbmc_4k | 4830 | 4335 | 0.9993 | 0.9993 | 0.9999 | 4231 | 0.9804 | 0.9991 | 0.9935 |
| t_4k_small | 4716 | 4494 | 0.9955 | 0.9993 | 0.9998 | 5004 | 0.8929 | 0.9993 | 0.9884 |
| placenta1 | 6285 | 5636 | 0.9963 | 0.9991 | 0.9999 | 7149 | 0.8741 | 1.0000 | 0.9983 |
| placenta2 | 2771 | 2101 | 0.9700 | 0.9990 | 0.9997 | 3530 | 0.7697 | 0.9999 | 0.9844 |
| placenta3 | 5373 | 4019 | 0.9988 | 0.9982 | 0.9997 | 4600 | 0.9833 | 0.9988 | 0.9987 |
| placenta4 | 5038 | 3906 | 0.9964 | 0.9984 | 0.9996 | 4532 | 0.9709 | 0.9991 | 0.9981 |
| placenta5 | 4064 | 2723 | 0.9927 | 0.9982 | 0.9994 | 3885 | 0.8976 | 0.9992 | 0.9972 |
| placenta6 | 3942 | 3289 | 0.9960 | 0.9991 | 0.9998 | 4238 | 0.9033 | 0.9998 | 0.9971 |
| 3659_colon | 3376 | 5649 | 0.5936 | 1.0000 | 0.9877 | 5655 | 0.5929 | 1.0000 | 0.9878 |

**Supplementary Table 4.** Global comparison statistics between dropkick, CellRanger_2 and EmptyDrops for 33 inDrop scRNA-seq datasets.

|  | dropkick label<br>total cells | CellRanger 2<br>total cells | CellRanger 2<br>sensitivity | CellRanger 2<br>specificity | CellRanger 2<br>AUC | EmptyDrops<br>total cells | EmptyDrops<br>sensitivity | EmptyDrops<br>specificity | EmptyDrops<br>AUC |
| --- | --- | --- | --- | --- | --- | --- | --- | --- | --- |
| 2771 S1 | 2222 | 2401 | 0.7380 | 0.9264 | 0.9263 | 2519 | 0.7594 | 0.9472 | 0.9590 |
| 2771 S2 | 1248 | 925 | 0.8465 | 0.9031 | 0.9237 | 1219 | 0.8515 | 0.9506 | 0.9634 |
| 2771 S3 | 1382 | 1422 | 0.6392 | 0.9643 | 0.9154 | 983 | 0.8688 | 0.9616 | 0.9771 |
| 3072 S1 | 962 | 1536 | 0.5137 | 0.9899 | 0.9619 | 1049 | 0.6616 | 0.9848 | 0.9788 |
| 3072 S2 | 2370 | 2239 | 0.6807 | 0.9434 | 0.9196 | 1571 | 0.8250 | 0.9322 | 0.9531 |
| 3072 S3 | 3112 | 2873 | 0.8190 | 0.9102 | 0.9331 | 3210 | 0.8059 | 0.9334 | 0.9485 |
| 3254 S1 | 1285 | 1356 | 0.8201 | 0.9536 | 0.9501 | 1235 | 0.8907 | 0.9521 | 0.9783 |
| 3777 S1 | 1066 | 828 | 0.9541 | 0.9082 | 0.9669 | 908 | 0.9361 | 0.9247 | 0.9735 |
| 3777 S2 | 927 | 1205 | 0.6929 | 0.9802 | 0.9435 | 1699 | 0.5221 | 0.9902 | 0.9712 |
| 3907 S1 | 2260 | 2016 | 0.8661 | 0.9788 | 0.9782 | 1948 | 0.9004 | 0.9792 | 0.9892 |
| 3907 S2 | 1828 | 2057 | 0.7321 | 0.9828 | 0.9652 | 1207 | 0.9205 | 0.9641 | 0.9840 |
| 4058 colon | 2186 | 2155 | 0.7940 | 0.9827 | 0.9788 | 1973 | 0.8246 | 0.9798 | 0.9790 |
| MPP 03A | 945 | 997 | 0.6861 | 0.9534 | 0.9224 | 914 | 0.8271 | 0.9663 | 0.9771 |
| MPP 04A | 702 | 766 | 0.8055 | 0.9365 | 0.9281 | 847 | 0.8040 | 0.9824 | 0.9842 |
| MPP 07A | 1384 | 1352 | 0.9098 | 0.9761 | 0.9837 | 1256 | 0.9594 | 0.9728 | 0.9895 |
| MPP 08A | 1473 | 1601 | 0.7077 | 0.9597 | 0.9290 | 1433 | 0.8130 | 0.9641 | 0.9692 |
| MPP 13A | 1815 | 1634 | 0.8782 | 0.9501 | 0.9563 | 1744 | 0.8893 | 0.9643 | 0.9744 |
| MPP 18A | 875 | 753 | 0.8539 | 0.9655 | 0.9558 | 661 | 0.9213 | 0.9612 | 0.9799 |
| MPP 19A | 1503 | 1477 | 0.8538 | 0.9732 | 0.9809 | 1416 | 0.8905 | 0.9734 | 0.9876 |
| MPP 21A | 2149 | 2096 | 0.8545 | 0.9522 | 0.9574 | 1518 | 0.9697 | 0.9193 | 0.9663 |
| MPP 48A | 811 | 287 | 0.9477 | 0.9389 | 0.9652 | 934 | 0.8084 | 0.9927 | 0.9931 |
| MPX 03A | 1228 | 1197 | 0.8797 | 0.9370 | 0.9471 | 1217 | 0.9260 | 0.9624 | 0.9872 |
| MPX 03B | 407 | 782 | 0.3964 | 0.9861 | 0.9343 | 493 | 0.5558 | 0.9818 | 0.9779 |
| MPX 04A | 744 | 975 | 0.6338 | 0.9900 | 0.9136 | 730 | 0.8274 | 0.9891 | 0.9878 |
| MPX 08A | 1277 | 1100 | 0.8236 | 0.9563 | 0.9599 | 906 | 0.9183 | 0.9492 | 0.9767 |
| MPX 08B | 947 | 962 | 0.8753 | 0.9889 | 0.9751 | 1050 | 0.8333 | 0.9923 | 0.9915 |
| MPX 09A | 549 | 507 | 0.7929 | 0.9655 | 0.9622 | 609 | 0.8144 | 0.9870 | 0.9857 |
| MPX 09B | 1462 | 1235 | 0.8899 | 0.9767 | 0.9785 | 1143 | 0.9283 | 0.9745 | 0.9850 |
| MPX 17A | 889 | 813 | 0.7282 | 0.9751 | 0.9638 | 1347 | 0.5457 | 0.9863 | 0.9709 |
| MPX 19A | 857 | 798 | 0.7406 | 0.9673 | 0.9630 | 865 | 0.7260 | 0.9715 | 0.9737 |
| MPX 19B | 2830 | 2753 | 0.8794 | 0.9493 | 0.9709 | 2769 | 0.9029 | 0.9586 | 0.9804 |
| MPX 21A | 1647 | 1629 | 0.8422 | 0.9660 | 0.9713 | 1622 | 0.8662 | 0.9700 | 0.9812 |
| MPX 21B | 1377 | 1287 | 0.8143 | 0.9541 | 0.9608 | 1020 | 0.9255 | 0.9426 | 0.9748 |

**Supplementary Table 5.**
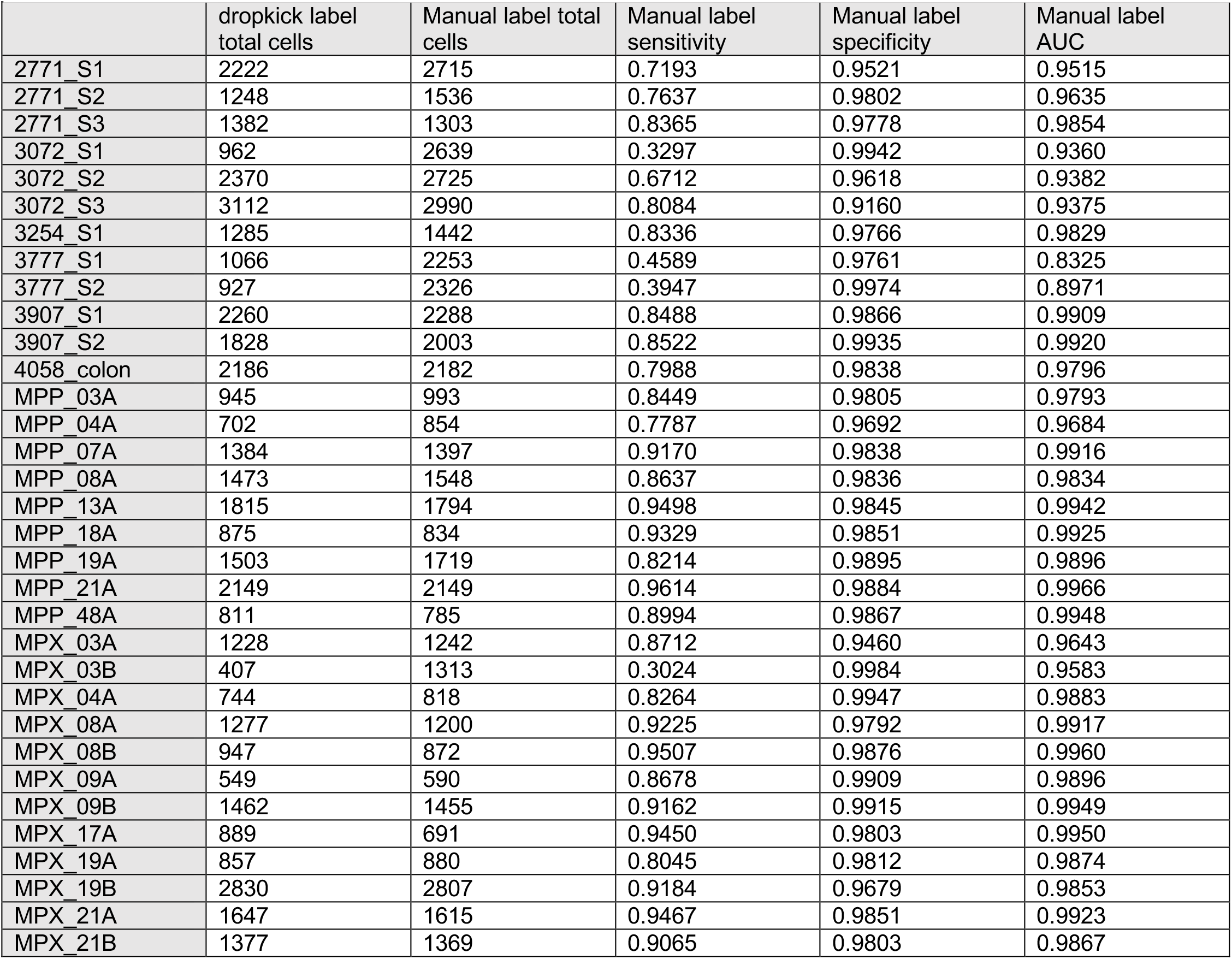
Global comparison statistics between dropkick and manual cell labelling for 33 inDrop scRNA-seq datasets.

|  | dropkick label<br>total cells | Manual label total<br>cells | Manual label<br>sensitivity | Manual label<br>specificity | Manual label<br>AUC |
| --- | --- | --- | --- | --- | --- |
| 2771 S1 | 2222 | 2715 | 0.7193 | 0.9521 | 0.9515 |
| 2771 S2 | 1248 | 1536 | 0.7637 | 0.9802 | 0.9635 |
| 2771 S3 | 1382 | 1303 | 0.8365 | 0.9778 | 0.9854 |
| 3072 S1 | 962 | 2639 | 0.3297 | 0.9942 | 0.9360 |
| 3072 S2 | 2370 | 2725 | 0.6712 | 0.9618 | 0.9382 |
| 3072 S3 | 3112 | 2990 | 0.8084 | 0.9160 | 0.9375 |
| 3254 S1 | 1285 | 1442 | 0.8336 | 0.9766 | 0.9829 |
| 3777 S1 | 1066 | 2253 | 0.4589 | 0.9761 | 0.8325 |
| 3777 S2 | 927 | 2326 | 0.3947 | 0.9974 | 0.8971 |
| 3907 S1 | 2260 | 2288 | 0.8488 | 0.9866 | 0.9909 |
| 3907 S2 | 1828 | 2003 | 0.8522 | 0.9935 | 0.9920 |
| 4058 colon | 2186 | 2182 | 0.7988 | 0.9838 | 0.9796 |
| MPP 03A | 945 | 993 | 0.8449 | 0.9805 | 0.9793 |
| MPP 04A | 702 | 854 | 0.7787 | 0.9692 | 0.9684 |
| MPP 07A | 1384 | 1397 | 0.9170 | 0.9838 | 0.9916 |
| MPP 08A | 1473 | 1548 | 0.8637 | 0.9836 | 0.9834 |
| MPP 13A | 1815 | 1794 | 0.9498 | 0.9845 | 0.9942 |
| MPP 18A | 875 | 834 | 0.9329 | 0.9851 | 0.9925 |
| MPP 19A | 1503 | 1719 | 0.8214 | 0.9895 | 0.9896 |
| MPP 21A | 2149 | 2149 | 0.9614 | 0.9884 | 0.9966 |
| MPP 48A | 811 | 785 | 0.8994 | 0.9867 | 0.9948 |
| MPX 03A | 1228 | 1242 | 0.8712 | 0.9460 | 0.9643 |
| MPX 03B | 407 | 1313 | 0.3024 | 0.9984 | 0.9583 |
| MPX 04A | 744 | 818 | 0.8264 | 0.9947 | 0.9883 |
| MPX 08A | 1277 | 1200 | 0.9225 | 0.9792 | 0.9917 |
| MPX 08B | 947 | 872 | 0.9507 | 0.9876 | 0.9960 |
| MPX 09A | 549 | 590 | 0.8678 | 0.9909 | 0.9896 |
| MPX 09B | 1462 | 1455 | 0.9162 | 0.9915 | 0.9949 |
| MPX 17A | 889 | 691 | 0.9450 | 0.9803 | 0.9950 |
| MPX 19A | 857 | 880 | 0.8045 | 0.9812 | 0.9874 |
| MPX 19B | 2830 | 2807 | 0.9184 | 0.9679 | 0.9853 |
| MPX 21A | 1647 | 1615 | 0.9467 | 0.9851 | 0.9923 |
| MPX 21B | 1377 | 1369 | 0.9065 | 0.9803 | 0.9867 |

**Supplementary Table 6.**
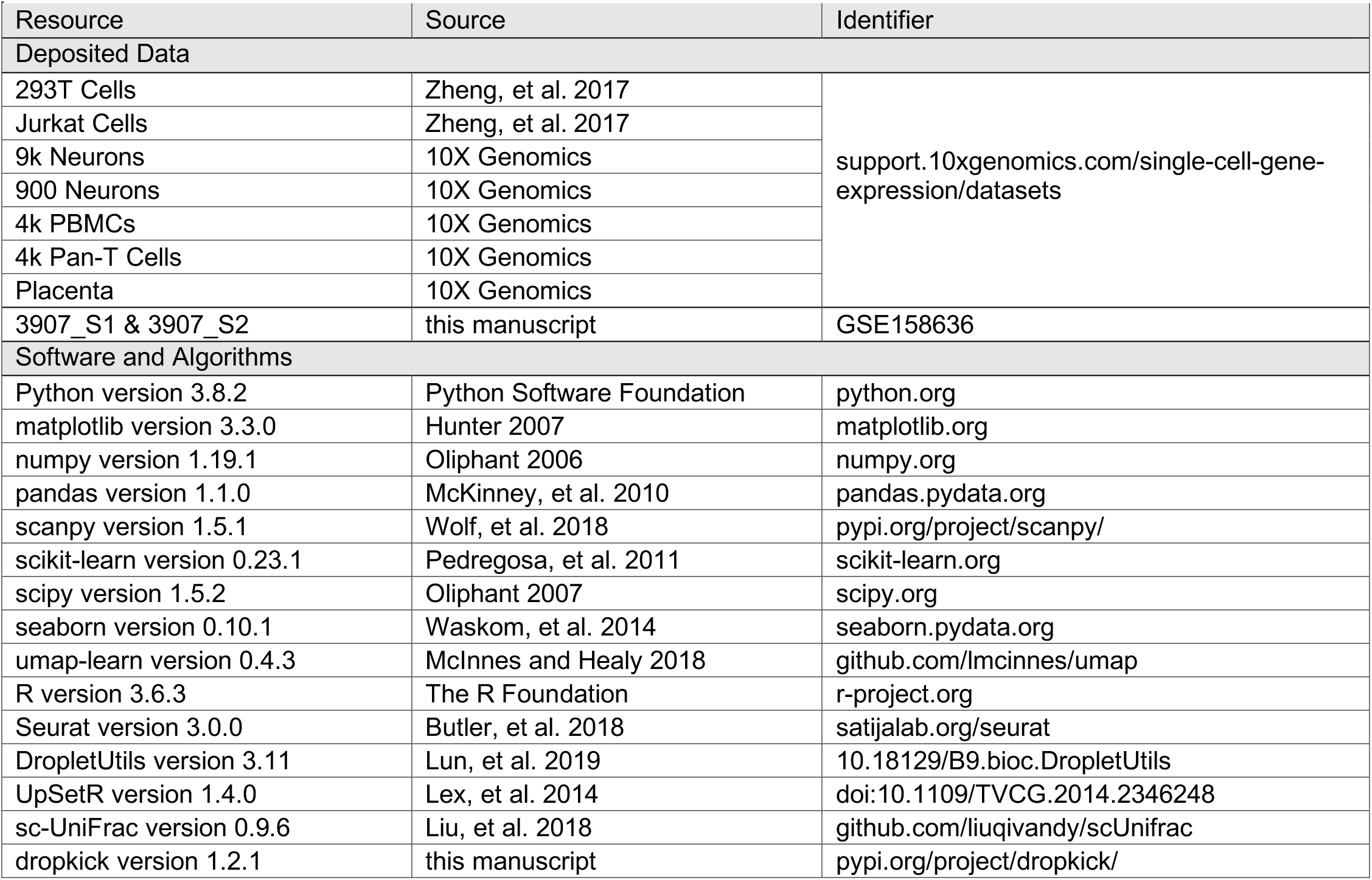
Code and data resources.

## Notes

https://github.com/KenLauLab/dropkick

### Competing Interest Statement

The authors have declared no competing interest.

